# Early-life stress exposure impacts the hippocampal synaptic proteome in a mouse model of Alzheimer’s disease: age- and pathology-dependent effects on mitochondrial proteins

**DOI:** 10.1101/2023.04.20.537660

**Authors:** Janssen M. Kotah, Mandy S.J. Kater, Lianne Hoeijmakers, Niek Brosens, Sylvie L. Lesuis, Roberta Tandari, Luca Marchetto, Ella Yusaf, August B. Smit, Paul J. Lucassen, Harm Krugers, Mark H.G. Verheijen, Aniko Korosi

**Author notes:** shared first authorship (Janssen M. Kotah, Mandy S.J. Kater). shared last authorship (Mark H.G. Verheijen, Aniko Korosi).

## Abstract

Epidemiological evidence indicates that early life stress (ES) exposure increases the risk for later-life diseases, such as Alzheimer’s disease (AD). Accordingly, we and others have shown that ES aggravates the development of, and response to, amyloid-beta (Aβ) pathology in animal models. Moreover, ES-exposed transgenic APP/PS1 mice display deficits in both cognitive flexibility and synaptic function. As the mechanisms behind these changes were unclear, we here investigated how exposure to ES, using the limited nesting and bedding model, affects the synaptic proteome across 2 different ages in both wildtype and APP/PS1 transgenic mice.

We found that, compared to wildtype mice, the hippocampal synaptosomes of APP/PS1 mice at an early pathological stage (4 months) showed a higher abundance of mitochondrial proteins and lower levels of proteins involved in actin dynamics. Interestingly, ES exposure in wildtype mice had similar effects on the level of mitochondrial and actin-related synaptosomal proteins at this age, whereas ES exposure had no additional effect on the synaptosomal proteome of early-stage APP/PS1 mice. Accordingly, ultrastructural analysis of the synapse using electron microscopy in a follow-up cohort showed fewer mitochondria in pre- and post-synaptic compartments of APP/PS1 and ES-exposed mice, respectively.

At a later pathological stage (10 months), the hippocampal synaptic proteome of APP/PS1 mice revealed an upregulation of proteins related to Aβ processing, that was accompanied by a downregulation of proteins related to postsynaptic receptor endocytosis. ES exposure no longer affected the synaptic proteome of wildtype animals by this age, whereas it affected the expression of astrocytic proteins involved in lipid metabolism in APP/PS1 mice. We confirmed a dysregulation of astrocyte protein expression in a separate cohort of 12-month-old mice, by immunostaining for the alpha subunit of the mitochondrial trifunctional protein and fatty acid synthase in astrocytes.

In conclusion, our data suggest that ES and amyloidosis share pathogenic pathways involving synaptic mitochondrial dysfunction and astrocytic lipid metabolism. These pathways might be underlying contributors to the long-term aggravation of the APP/PS1 phenotype by ES, as well as to the ES-associated risk for AD progression.

These data are publicly accessible online as a web app via https://amsterdamstudygroup.shinyapps.io/ES_Synaptosome_Proteomics_Visualizer/.

## Background

Despite advances in our understanding of the role of genetics in Alzheimer’s disease (AD) (1,2), recent evidence emphasizes the important role of environmental (i.e., non-genetic) factors in its etiology (3) including (chronic) stress (4–6). In particular, there is emerging clinical and epidemiological evidence that a history of early-life stress (ES) might increase the risk to develop cognitive decline (7–9), AD, and other dementias (9–18).

This association has been further substantiated in transgenic APPswe/PS1de9 mice, a mouse model of amyloidosis, which overexpress mutated versions of the human amyloid precursor protein and presenilin-1 (19,20). These mutations result in amyloid-beta (Aβ) plaques by 6 months of age (19,20), accompanied by altered cognition (21–23), synaptic protein expression (24), and neuroinflammation (22,25), among other phenotypes, as described in (26). When APP/PS1 mice are exposed to ES, via e.g. the limited bedding and nesting (LBN) model (27,28), we and others have reported age-dependent alterations in plaque load (29) and neuroinflammation (29,30), as well as deficits in cognitive flexibility (31), fear learning (32,33), postsynaptic receptor protein expression (33), and long term potentiation (34).

The synapse is a prominently affected substrate in AD patients, who exhibit synaptic loss, especially in the hippocampus (35,36). This synaptic loss is thought to occur early on and to underlie the cognitive symptoms of the disease (37), leading to a view of AD as a synaptic disorder (38,39). Synaptic deficits precede neurodegenerative features of AD, are present in early and prodromal cases (40,41), and have been suggested in animal models to increase as Aβ pathology accumulates (24,42). Therefore, synapse (dys)function might be a key substrate impacted both in early AD as well as a result of ES exposure.

We therefore set out to study how the hippocampal synaptic proteome is affected in APP/PS1 mice during early, pre-plaque stages of pathology, and how these changes compare to more advanced pathological stages. We further assessed how prior ES exposure during the first postnatal week (via the LBN model) might further modulate these genetically encoded alterations. We studied this using label free mass spectrometry in APP/PS1 mice and their wildtype (WT) littermates at 4 and 10 months of age (4 moa/10 moa), representing stages with low and high Aβ pathology, respectively (29). The hippocampi used in this study were derived from a cohort of APP/PS1 mice from which we previously reported the effects of ES on cognition (43), neurogenesis (43), and neuroimmune (i.e., microglial (29) and astrocytic (30)) profiles.

We found similar changes in synaptosomal proteins associated with mitochondria and actin dynamics in both non-stressed APP/PS1 and control ES mice at 4 moa, with no further alterations in ES-exposed APP/PS1 mice. At 10 moa, Aβ-associated alterations were pronounced in APP/PS1 brains, with previous ES-exposure also further altering (mitochondrial) lipid metabolism in APP/PS1 mice. These data reveal changes not only in neuronal proteins but also in proteins that are highly expressed by microglia and astrocytes. We then characterized ultrastructural mitochondrial alterations by performing electron microscopy in mice at 4 moa, and validated our main findings via immunostaining, showing that alterations in astrocytic lipid proteins in ES-exposed APP/PS1 mice persisted until 12 months.

## Materials and Methods

### Animal use

For this study, we used male bigenic hemizygous APPswe/PS1dE9 (TG) mice on a C57Bl/6J background and their wildtype (WT) littermates. To investigate how synaptic protein expression is altered by Aβ overexpression, we compared WT and TG mice sacrificed at 4 months of age (moa, Cohort 1: WT n=7, TG n=7). Next, to test how early-life stress (ES) influences the Aβ-induced synaptic protein alterations, we exposed WT and TG mice exposed to control (CTL) or ES conditions during the first postnatal week and sacrificed them at either at an early pathological stage (4 moa, Cohort 2: WT-CTL n=5, WT-ES n=4, TG-CTL n=4, TG-ES n=4) or at an advanced pathological stage (10 months of age/10 moa, Cohort 3: WT-CTL n=5, WT-ES n=9, TG-CTL n=5, TG-ES n=6), respectively.

Two more cohorts were generated to follow up on the proteomics data. Cohort 4 consisted of ES-exposed WT and TG littermates that were sacrificed at 4 moa to investigate mitochondria using electron microscopy (WT-CTL: n=7, WT-ES: n=5, TG-CTL: n=5, TG-ES: n=6). Cohort 5 consisted of CTL or ES-exposed TG mice sacrificed at 12-months and was used to validate two proteins that were found to be differentially expressed in the 10 moa proteomics data (TG-CTL: n=8, TG-ES: n=6).

Mice were bred in-house and kept under standard housing conditions (temperature 20–22 °C, 40–60% humidity level, chow/water ad libitum, 12/12 h light/dark schedule). All experiments were approved by the Animal Experiment Committee of the Vrije University Amsterdam and University of Amsterdam.

### Early-life stress paradigm

Mice used in ES experiments were bred and randomly assigned to control or ES groups at postnatal day (PND) 2 as previously described (28). Briefly, CTL litters were housed in standard amounts of sawdust, along with a 5x5cm piece of nesting material (Tecnilab-BMI, Someren, The Netherlands). ES nests were housed on a fine-gauge stainless steel mesh on top of 1/3 of the regular amount of CTL bedding material, along with half of a 2.5x5 cm square of nesting material. All nests were covered with filter tops. Nests were transferred to standard cage conditions at PND9 and allowed to age until used in our experiments.

### Synaptosome proteomics

#### Tissue collection and synaptosome isolation

To study how Aβ pathology affects the hippocampal synaptosome, mice from Cohorts 1, 2, and 3 were sacrificed via rapid decapitation in the morning. The hippocampi of these mice were dissected, snap frozen on dry ice, and stored at -80°C until further use.

Synaptosomes were isolated on a discontinuous sucrose gradient as described previously (44,45). In short, homogenization buffer (0.32 M sucrose, 5 mM HEPES, in PBS pH=7.4, with protease inhibitor cocktail [Roche]) was added to hippocampi samples, after which they were mechanically homogenized by a Dounce homogenizer (12 strokes, 900 rpm). Samples were centrifuged at 1000 x *g* for 10 min and supernatant was collected. This was layered on a 0.85/1.2 M sucrose gradient and centrifuged at 30,000 x *g* for 2h. Synaptosomes were collected from the interface, which were further diluted with 5 ml homogenization buffer and centrifuged at 20,000 x *g* for 30 min to obtain a synaptosome pellet. All steps were performed on ice.

#### FASP in-solution digestion of proteins

Samples were digested by filter-aided sample preparation (FASP) as previously described (45). Ten µg of synaptosomes from each sample was incubated with 75 µl reducing agent (2% SDS, 100 mM TRIS, 1.33 mM TCEP) at 55 °C for 1h at 900 rpm and subsequently incubated with 500 mM MMTS for 30 min at room temperature (RT). The samples were transferred to YM-30 filters (Microcon, Millipore) and a washing solution (200 µl 8 M urea in 100 mM TRIS (pH=8.8)) was added to wash five times by spinning at 14,000 x *g* for 15 min. Next, the samples were washed with 50 mM NH_4_HCO_3_ for four more times. Samples were incubated overnight at 37 °C with 100 µl of Trypsin (0.6 grams in 100 µl 50 mM NH_4_HCO_3_). Digested peptides were eluted from the filter with 50 mM NH_4_HCO_3_. The samples were dried using a SpeedVac and stored at -20 °C.

#### Mass spectrometry-based analysis

Samples were loaded onto an Ultimate 3000 LC system (Dionex, Thermo Scientific) as described previously (24,44–46). Spectronaut 14 (Biognosys) was used for data analysis of the raw files. The spectral library was created with crude hippocampal synaptosomes containing spiked-in peptides (Biognosys), analysed with a TripleTOF 5600 in data-dependent acquisition mode. The obtained library was searched against mouse proteome (UP000000589_10090.fasta and UP000000589_10090.addition.fasta) in MaxQuant Software (version 1.3.0.5). Data quality control and statistical analysis were performed by using the downstream analysis pipeline for quantitative proteomics (MS-DAP version 0.2.6.3 (47)). Outliers were removed in case of large within or between sample variations observed by deviating distribution plots, or in case of disturbed protein detection observed by altered retention time plots.

#### Data analysis

Peptide abundance values were normalized and the MSqRob algorithm was used for peptide-level statistical analysis. The threshold for significance was set at FDR<0.05 and Log_2_ fold change cut off at -0.1/0.1. Venn diagrams were created using the ggVenndiagram package on R, v1.1.0 (48). To gain insight into whether specific cell-types are more altered based on the differentially expressed proteins, we performed expression weighted cell-type enrichment (EWCE) analysis (49), using a hippocampal single cell RNAseq dataset (50).

Functional insights for differentially expressed proteins were obtained by separately analyzing significantly dysregulated up- and down-regulated proteins with gene ontology (GO, based on biological processes [BP] and cellular components [CC]) using gProfiler2 (51), as well as SynGO, a curated resource of synaptic proteins (52). Because a majority of our significantly dysregulated proteins were not annotated in SynGO, we chose to use GO as our main downstream analysis, and to analyze our proteins in SynGO only when we obtained neuronal enrichment from EWCE analyses. GO and enrichment analyses were only performed when there were at least 5 differentially expressed proteins, and GO analyses that yielded >10 significantly overrepresented terms (FDR < 0.05) were also semantically clustered using RRVGO to reduce redundancy (53). Mitochondrial proteins and pathway analyses were done by importing annotations from the MitoCarta3.0 database (54) into gProfiler2. All data visualizations and statistics were performed in RStudio v1.4.1717 (55).

### Ultrastructural analysis of mitochondria

#### Tissue preparation

To further understand what the identified differences in mitochondrial proteins in TG and ES mice at 4 moa mean for mitochondrial structure, we subsequently studied these at the ultrastructural level via electron microscopical analysis. Mice at 4 moa (Cohort 4) were sacrificed via transcardial perfusion with ice-cold 4% paraformaldehyde (PFA) in phosphate buffered saline (PBS, pH = 7.4) under anesthesia by 120mg/kg Euthasol. Whole brains were dissected and kept in 4% PFA for 24 h after which the solution was replaced for 30% sucrose for cryopreservation of the tissue. Brain tissue was stored at -80 °C until further processed.

Fifty µm thin coronal sections of the hippocampus were made on a sliding microtome. Contrasting of the sections was realized using a solution of 1% osmium and 1% ruthenium. The slices were then exposed to increasing ethanol concentrations (30%, 50%, 70%, 90%, 96% and 100%) and finally propylene oxide for dehydration. This was followed by embedding the sections in epoxy resin and polymerization for 72 h at 65 °C. Ultra-thin sections of 90 nm of the dorsal CA1 hippocampus were cut on an ultra-microtome (Reichert-Jung, Ultracut E). Finally, post-contrast was realized with uranyl acetate and lead citrate in an ultra-stainer (Leica EM AC20).

#### Electron microscopy imaging and analysis

The grids were examined with a JEOL JEM 1011 electron microscope at 50.000x magnification. Pictures were taken with an Olympus Moreda 11-MP camera and iTEM software (Olympus). A total of 70 pictures of randomly selected locations within a section of dorsal CA1 hippocampus (around Bregma point -1.6) were collected in which at least one morphologically intact mitochondrion was identified. Mitochondria within peri-synaptic structures were counted and classified as belonging to either pre-synapse (based on presence of vesicles), post-synapse (based on presence of postsynaptic densities), dendritic spines (based on elongated form), or astrocytes (based on clear, electron deficient cytoplasm). Morphological features (Perimeter, Area, Circularity) of whole mitochondria (i.e., those not on the edge of the image) were measured using ImageJ. Outliers in the morphological measurements were removed using the 1.5* interquartile range method. Statistical analyses were done by creating a mixed model to assess the effects of genotype, early-life condition, and their interaction, while correcting for the nesting effect of multiple mitochondrial measures being obtained from the same mice. Post-hoc analyses were conducted correcting using Tukey’s method.

### Immunofluorescence

To validate ES-induced modulation of the hippocampal synaptic proteome in APP/PS1 mice at advanced stages of pathology, we used two parallel series of 40µm thick coronal brain slices from a previously described cohort of 12-month old APP/PS1 mice exposed to ES (32) (Cohort 5). These mice had been sacrificed via rapid decapitation within the first two hours of the light phase, and brains were immersion-fixated in 4% paraformaldehyde overnight, then stored in 0.1M Phosphate Buffer with 0.01% Na-Azide until slicing and stored in antifreeze at -20°C until use.

We immuno-stained against two proteins involved in lipid metabolism, i.e. Hydroxyacyl-CoA Dehydrogenase Trifunctional Multienzyme Complex Subunit Alpha (Hadha, rabbit polyclonal Abcam, ab54477, 1:250) and Fatty Acid Synthase (Fasn, rabbit polyclonal, Abcam ab22759, 1:500). Both were co-stained with Vimentin (chicken polyclonal, EMD Millipore 5733, 1:3000 for Hadha; 1:2000 for Fasn) to localize the signals to astrocytes.

Hadha immunostaining was performed on pre-mounted slices after 15 minutes of antigen retrieval at 100°C in citrate buffer (pH 6.0), while Fasn was performed free floating at RT. Both were blocked for 1h in a 0.05M TBS mix with 5% NGS and 0.3% Triton (pH 7.6), then incubated with primary antibodies for 1h at RT, then overnight at 4°C. Sections were incubated the next day with secondary antibodies (goat anti-chicken-A488, goat-anti-rabbit-A568, and goat-anti-mouse-A647, Invitrogen, 1:800) for 2h at RT. Slices were washed with x 0.05M TBS (pH7.6) between steps. We used 12 brains (TG-CTL: n=7, TG-ES: n=5) for Hadha staining and 14 brains (TG-CTL: n=8, TG-ES: n=6) for Fasn staining.

### Confocal microscopy and image analysis

We imaged 1µm-step Z-stacks (total Z range of 9-10µm) at 40x magnification using a Nikon A1 confocal microscope. Images were taken from six sections along the dorsoventral axis of the hippocampus to obtain an even representation. Each image was digitally stitched into one composite image such that all hippocampal subregions (Stratrum Oriens in the Cornu Ammonis [CA] to the Hilus in the dentate gyrus) were visible, with dorsal sections being 2 images wide and 4 images tall and ventral sections being 4 or 5 images wide and 3 images tall.

Images were analyzed with FIJI (v1.53q) by first drawing regions of interest to define the CA and dentate gyrus regions. Per slice, binary masks of Vimentin signal were generated using an automated threshold followed by particle analysis to reduce noise. The area of Vimentin signal, as well the automated thresholded area of the Hadha or Fasn signal were measured, with coverage being defined as (Hadha or Fasn area/Vimentin area). After testing for outliers using the 1.5*interquartile range method, data were analyzed using the student’s t-test.

### Visualizing trajectories of proteomic alterations from 4 moa to 10 moa

To visualize the trajectory of TG-induced effects at the synapse, we adapted a ΔLog2FC approach to visualize temporal dynamics of changes in synaptosomal proteomes (24). We performed this analysis in differentially expressed mitochondrial proteins when comparing either WT-CTL vs TG-CTL mice or TG-CTL vs TG-ES mice. For each protein, we calculated a ratio between their Log2FC across ages (Log2FC_10moa_/Log2FC_4moa,_ or ΔLog2FC). Using a threshold of ±0.05, we then stratified the differentially expressed proteins by whether their ΔLog2FC ratio increased (i.e., ΔLog2FC > 0.05), decreased (i.e., ΔLog2FC < -0.05), or was not affected (-0.05 < ΔLog2FC < 0.05) between 4 moa and 10 moa.

## Results

### Detection of synapse associated proteins from isolated synaptosomes

We confirmed the presence of bona fide synaptic proteins within our isolated hippocampal synaptosomes by analyzing gene set enrichment of total detected proteins against a background of brain expressed genes in SynGO, a curated resource of 1113 synaptic proteins (52). Out of around 3000 proteins detected per contrast, at least 674 proteins per contrast mapped to SynGO (**fig. S1A**), encompassing the spectrum of annotated cellular components and biological processes terms in the database (**fig. S1B**). We also found an overall overrepresentation of synaptic ontology terms in our detected proteins by performing GO analysis against the ‘brain background geneset’ used in SynGO (**fig. S1C**). This provided us with confidence in the success of our synaptosomal enrichment to proceed with downstream analyses.

### Early Aβ pathology alters hippocampal synaptic proteins involved in mitochondria and actin dynamics

To identify possible synaptic alterations induced by the transgenic APP/PS1 genotype (TG) at early stages of Aβ pathology, we characterized the proteome in hippocampal synaptosomes from wildtype (WT) and TG mice at 4 months of age (4 moa; Cohort 1, **fig. 1A**). Because the cohort generated to investigate early-life stress (ES) effects on TG mice also included WT and TG that were not exposed to ES (Cohort 2, **fig. 1B**), we present them here together. We found a similar number of differentially expressed proteins in both cohorts (Cohort 1: 14 up, 23 down; Cohort 2: 17 up, 23 down, **fig. 1C-E, table S1, S2**), of which two proteins (Abi1, Wasf) were downregulated in both cohorts.

**Figure 1.**
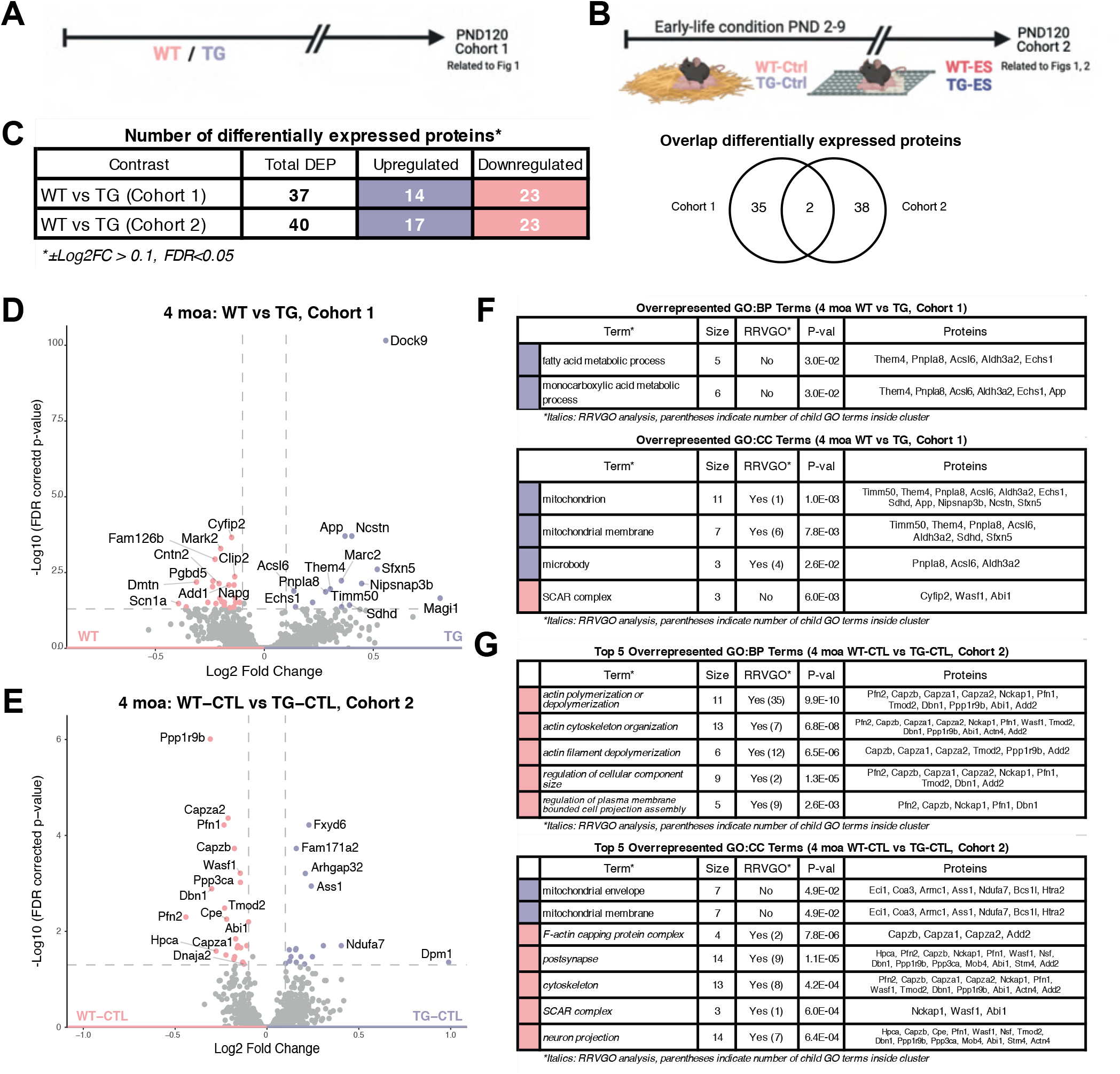
Synaptosomal upregulation of mitochondrial proteins and downregulation of actin-dynamics proteins in 4 moa APP/PS1 mice. (A) Experimental design in analyzing the hippocampal synaptosomal proteome at four months of age (4 moa). (B) Table summarizing differentially expressed proteins between two cohorts of wildtype (WT) and transgenic APP/PS1 (TG) mice. Only two proteins (Abi1, Wasf) were significantly downregulated in both cohorts. (C-D) Volcano plots showing differentially expressed proteins (±Log2FC > 0.1, FDR> 0.05) in cohorts 1 (C) and 2 (D). (E-F) Overview of overrepresented biological processes (BP) and cellular component (CC) terms based on gene ontology (GO) analyses of up- (light blue) and downregulated (light red) proteins in Cohorts 1 (E) and 2 (F). Italicized terms are ‘parent’ GO terms after clustering by semantic similarity using RRVGO.

Overrepresented GO terms, based on differentially expressed proteins in both cohorts, were related to the same processes, i.e., mitochondria and actin dynamics (**fig. 1F, 1G, table S3**), but notably involved different sets of proteins. Both cohorts had overrepresentation of terms related to the mitochondrial membrane (Timm50, Them4, Pnpla8, Acsl6, Aldh3a2, Sdhd, Sfxn5) in upregulated proteins, which were also overrepresented for fatty acid and monocarboxylic acid metabolism (Acsl6, Aldh3a2, App, Echs1, Pnpla8, Them4) in Cohort 1. This mitochondrial alteration was also evident when comparing the upregulated proteins with a curated database of mitochondrial proteins (MitoCarta3.0 (54)), with 9 such proteins in Cohort 1 (Acsl6, Aldh3a2, Echs1, Nipsnap3b, Pnpla8, Sdhd, Sfxn5, Them4, Timm50) and 7 in Cohort 2 (Bcs1l, Coa3, Eci1, Etfa, Htra2, Lactb, Ndufa7, Prkaca).

We also found in both cohorts a downregulation of proteins (*Cohort 1*: Abi1, Cyfip2, Wasf1; *Cohort 2*: Abi1, Nckap1, Wasf) involved in the SCAR complex, which plays a role in actin dynamics and cytoskeletal regulation (56), which notably included the two shared downregulated proteins (Abi1, Wasf). The downregulation of actin-related proteins in synaptosomes from TG mice was particularly evident in Cohort 2 (**fig. 1F**), wherein actin-related biological processes (e.g., polymerization/depolymerization: Abi1, Add2, Capza1, Capza2, Capzb, Dbn1, Nckap1, Pfn1, Pfn2, Ppp1r9b, Tmod2) were downregulated. In terms of cellular component annotations, these proteins were localized to the cytoskeleton, as well as in the post-synapse (**fig. 1G**, **table S3**).

To assess whether the list of up- and downregulated proteins in both cohorts reflected cell-type specific TG effects, we performed expression-weighted cell-type enrichment (EWCE) analysis (49) using a published single-cell gene expression database from the mouse hippocampus (50). TG synaptosomes from Cohort 1 exhibited astrocytic enrichment in the upregulated proteins and neuronal enrichment in downregulated proteins (**fig. S2A**). In Cohort 2, we did not detect cell type enrichment in upregulated proteins, while downregulated proteins were enriched for neurons (**fig. S2B**). Because of the neuronal annotation of downregulated proteins in both cohorts, we then further characterized their function using SynGO, and found downregulated proteins to be involved in postsynaptic processes, specifically with respect to actin-related processes (**fig S2C-D**).

### Early-life stress downregulates actin dynamics and upregulates astrocytic proteins in hippocampal synaptosomes from WT mice at 4 moa

We then investigated how early-life stress (ES) exposure affected the synaptic proteome in 4-month-old WT and TG mice (**fig. 2A**). ES exposure in WT animals resulted in 126 differentially expressed synaptosomal proteins (84 up, 42 down, **fig. 2B, table S4**). Based on GO analyses, a protein group enriched in WT-ES synaptosomes contains e.g. Atp1a2, Atp1b2, Gnai2, Slc1a3, Slc8a1, i.e. ATPase subunits as well as Glutamate and GABA transporters, and were associated with cellular component terms such as the GTPase complex, and endoplasmic reticulum of the cell (**fig. S2E**, **table S3**). We also found evidence for mitochondrial involvement in the upregulated proteins contrast, with an overrepresentation of mitochondria-associated membrane of the endoplasmic reticulum (Bcap31, Canx, Tmx2, **table S3**). The 126 upregulated proteins were revealed by EWCE analysis to have enriched astrocytic annotations (**fig 2C**). In line with this, some of these upregulated proteins, e.g. Atp1a2 (57), Slc1a2 (57), Slc1a3 (57,58), and Slc6a11 (59), have been described in literature to be specific for astrocytic proteins enriched around the synapse. GO analysis of downregulated proteins revealed overrepresentation of biological processes terms such as actin cytoskeletal dynamics (via Abi2, Actn4, Add2, Bin1, Capza1, Capza2, Capzb, Cfl1, Coro1c, Dbn1, Pfn2, Ppm1e, Ppp1r9b, Tmod2, Twf2), and cellular component terms such axonal growth cones (via Cfl1, Dbn1, Fkbp4, Hsp90ab1, Ppp1r9b, Tmod2, Twf2, **fig. S2F**, **table S3)**.

**Figure 2.**
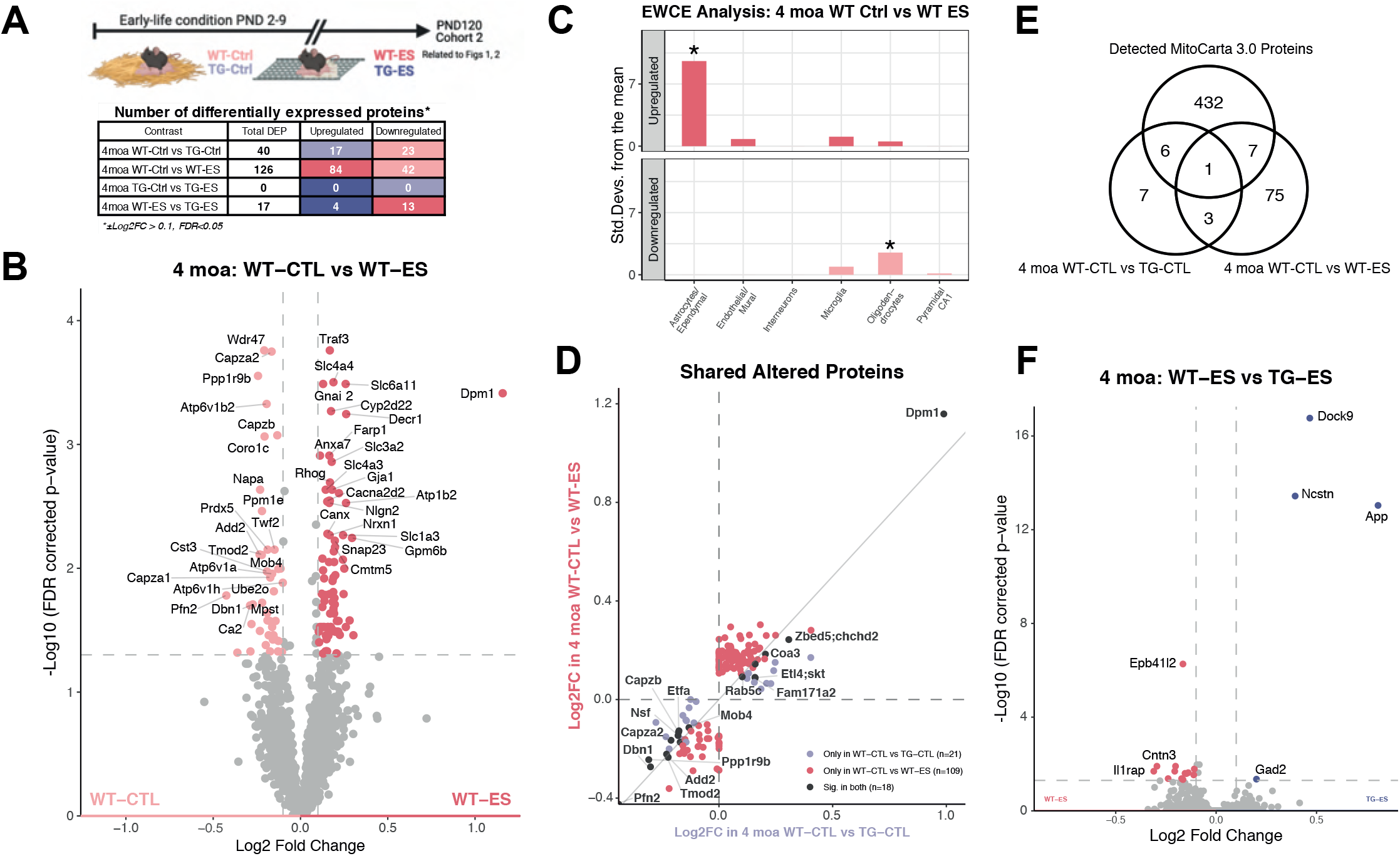
Early-life stress (ES) exposure upregulates astrocytic mitochondrial proteins and downregulates actin-dynamics related proteins. (A) Experimental design within cohort 2, consisting of wildtype (WT) and transgenic APP/PS1 (TG) mice exposed to control (CTL) or early-life stress (ES) at 4 months of age (4 moa), and a summary of differentially expressed proteins across contrasts. (B) Volcano plot showing differentially expressed proteins (±Log2FC > 0.1, FDR> 0.05) in synapses from ES-exposed WT mice. (C) Expression weighted celltype enrichment (EWCE) analysis shows enrichment of astrocytic annotations in upregulated proteins and oligodendrocytic annotations in downregulated proteins. (D) Overlap of differentially expressed proteins when comparing proteomic changes in synaptosomes from TG (light blue) and ES (dark red) mice. (E) Upregulated proteins in synaptosomes from TG-CTL and WT-ES mice are mitochondrial, as determined by the MitoCarta database. (F) Volcano plot showing differentially expressed proteins (±Log2FC > 0.1, FDR> 0.05) when comparing synapses from ES-exposed WT and TG mice.

### Possible shared effects between ES exposure and APP/PS1 genotype on the hippocampal synaptic proteome at 4 moa

ES exposure did not lead to significant changes in synaptosomal protein expression within TG mice at this age (**table S5**). This stands in contrast to the detected alterations to synaptosomes when comparing either TG-CTL or WT-ES mice to WT-CTL mice, suggesting a degree of convergence between ES and TG effects at this age. To explore this, we investigated the 18 proteins (7 up, 11 down) that were differentially expressed in the synaptosomes of both TG-CTL or WT-ES mice (**fig. 2D, table S7**). While the group of upregulated proteins did not show any overrepresentation of GO terms, the downregulated proteins were associated with actin-related biological processes (e.g., actin polymerization or depolymerization: Add2, Capza1, Capza2, Capzb, Dbn1, Pfn2, Ppp1r9b, Tmod2) and were overrepresented for cellular component terms such as the postsynapse (Add2, Capzb, Dbn1, Mob4, Nsf, Pfn2, Ppp1r9b) and dendritic spines (Capzb, Dbn1, Mob4, Ppp1r9b, **table S8**).

Beyond shared alterations to actin-related processes, inspection of the shared upregulated proteins also suggested mitochondrial changes in both ES and TG synaptosomes. This is evidenced firstly by the reported mitochondrial expression of Dpm1 (60), the protein with the highest log2 fold change value in TG-CTL (**fig. 1D**) and WT-ES (**fig. 2B**) synaptosomes. Several shared upregulated proteins are also either expressed in mitochondria (e.g. Coa3 (61), Chchd2 (62,63)), or have been reported to interact with mitochondria and to regulate their function (e.g. Rab5c (64,65), Etl4-Skt (66), Slc4a3 (67)). Lastly, several upregulated proteins in synaptosomes isolated from WT-ES mice overlapped with MitoCarta (Acaa2, Acad8, Acadm, Coa3, Decr1, Glud1, Idh2, Etfa, Mpst, Ndufa3, Prdx5, **fig. 2E**). similar to the proteins identified when comparing TG-CTL vs WT-CTL synaptosomes.

Next to the overlap between ES and TG effects on synaptosomal protein expression, comparison of ES-exposed WT and TG mice revealed an additional 17 proteins as a result of Aβ overexpression (4 up, 13 down, **fig. 2F, table S6**). Upregulated proteins here are involved in Aβ processing (App, Dock9, Ncstn) and GABA biosynthesis (Gad2), while SynGO:BP and GO:CC analyses revealed downregulated proteins associated with postsynaptic membrane receptor levels (Ctnnd2, Agap3, Snap23, **fig. S2G**) and the hippocampal CA3 mossy fiber synapse (Adcy1, Shank2, Syt7, **table S3)**, respectively.

### Ultrastructural analyses of synaptic mitochondria at 4 moa reveal a distinct loss of mitochondrial numbers in the hippocampus of both APP/PS1 and ES-exposed mice

Based on the recurrence of mitochondrial terms in our proteomics data, we hypothesized that this organelle might be a substrate through which Aβ overexpression, ES-exposure, or both alter the hippocampal synapse. We explored this in a follow-up cohort of 4 moa WT and TG mice exposed to ES to study their mitochondria at the ultrastructural level (**fig. 3A).** We determined the number of mitochondria and assessed mitochondrial morphology at the ultrastructural level within the presynapse, postsynapse, dendrites, and astrocytes in the CA1 subregion of the hippocampus of these mice (**fig. 3B**).

**Figure 3.**
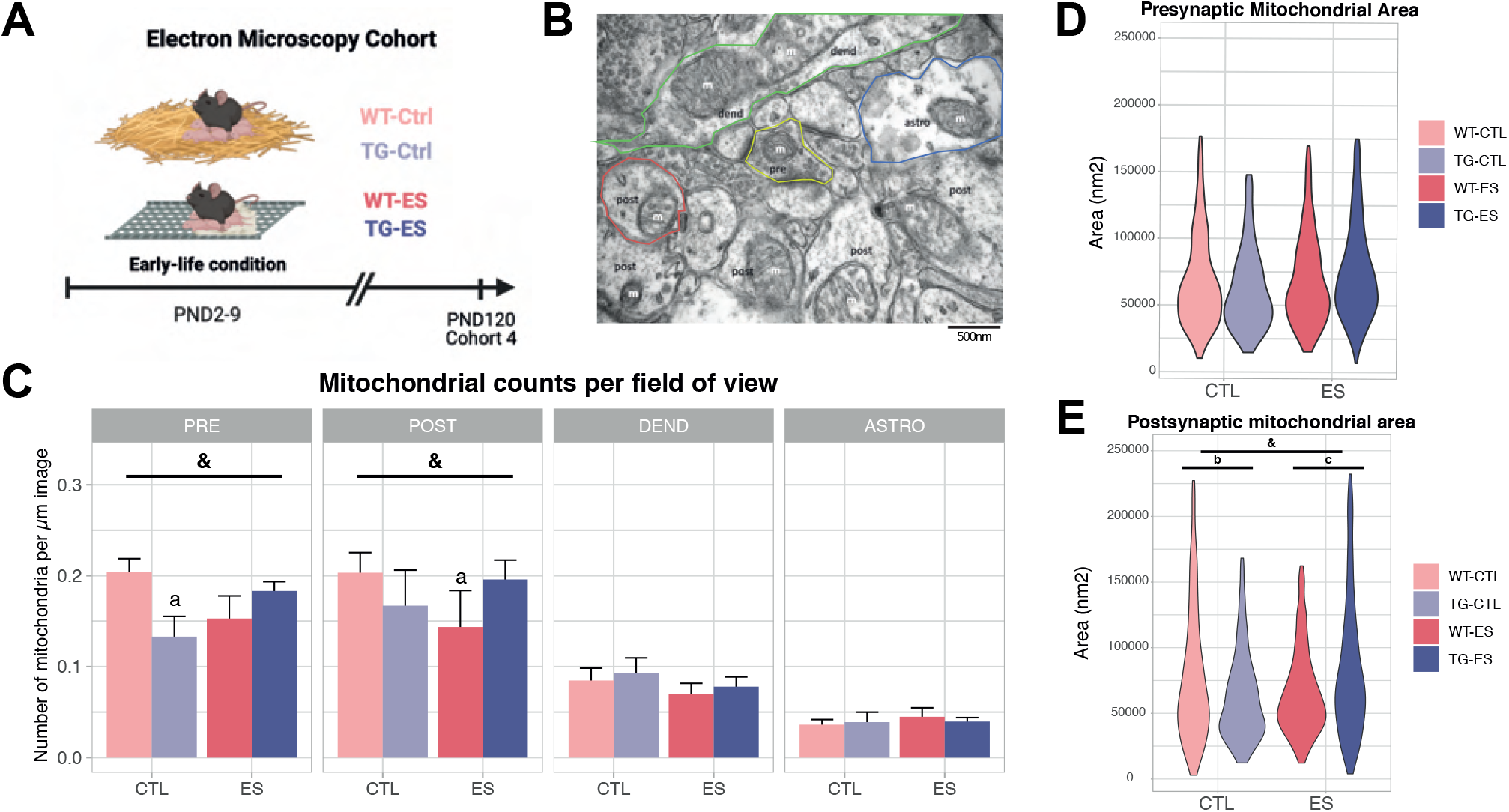
Investigating ultrastructural alterations to synaptic mitochondria in ES-exposed WT and TG mice. (A) Overview of cohort 4, consisting of wildtype (WT) and transgenic APP/PS1 (TG) mice exposed to control (CTL) or early-life stress (ES) at 4 months of age (4 moa), which was generated for electron microscopy (EM) analysis of mitochondria. (B) Representative EM image analyzed. Mitochondria (m) at the CA1 region of the hippocampus were counted and traced across 70 images per animal to analyze their morphology. Analysis was done separately in sub-synaptic structures, as labeled and outlined in the image: presynapse (yellow), postsynapse (red), astrocyte (blue), dendritic compartment (green). (C) Interaction of TG and ES effects on mitochondrial counts at the pre- and post-synapse, but not in dendrites or astrocytes. (D) Alterations to presynaptic mitochondrial counts are not accompanied by alterations to mitochondrial size. (E) Interaction of TG and ES effects on mitochondrial area at the postsynapse. &, interaction effect, p>0.05; post-hoc effects: a, different from WT-CTL, p>0.05; b, difference between WT-CTL and TG-CTL, p>0.05; c, difference between WT-ES and TG-ES, p>0.05. Analyses performed using a mixed linear model, WT-CTL: n=7, WT-ES: n=5, TG-CTL: n=5, TG-ES: n=6.

While there was no overall TG or ES effect on total number of mitochondria (**fig. S3A**), we found interaction effects of both factors on the number of mitochondria in the pre-synapse (condition: F(1,19)=0.0218, p=0.8842; genotype: F(1,19)=2.1204, p=0.1617; interaction: F(1,19)=12.001, p=0.0026) and post-synapse (condition: F(1,16)=2.5384, p=0.1307; genotype: F(1,16)=0.0555, p=0.8168; interaction: F(1,16)=11.8658, p=0.0033, **fig. 3C**). Mitochondria within astrocytes or dendritic structures were not affected. Using pairwise post-hoc tests, we found the pre-synaptic effect to be due to a decrease in mitochondria in TG-CTL compared to WT-CTL synapses (p=0.0109), whereas the postsynaptic interaction effect was driven by a decrease in the number of mitochondria in WT-ES compared to WT-CTL synapses (p=0.0171).

The effects on mitochondrial numbers were not accompanied by alterations in mitochondrial surface area at the pre-synapse (**fig. 3D**), although there was an interaction between TG and ES effects on the postsynaptic mitochondrial area (**fig. 3E**, condition: t(20)=-1.7961, p=0.0876; genotype: t(20)=-2.2274, p=0.0376; interaction: t(20)=3.3099, p=0.0035). Post-hoc analyses reveal that this interaction is explained by significant but opposite effects of genotype in CTL (t(20)=2.227, p=0.0376) and ES (t(20)=-2.452, p=0.0235) synapses. Mitochondrial area in other regions, as well as perimeter and circularity, were not significantly altered by either experimental variable (**fig. S3B-D**).

### At 10 months of age, APP/PS1 synaptosomes are depleted in proteins involved in presynaptic vesicle release and postsynaptic receptor endocytosis

We then characterized the effects of TG genotype and prior ES exposure on the hippocampal synaptosomal proteome in 10-month-old (10 moa) mice (**fig. 4A**). Synaptosomes isolated from TG-CTL mice showed large protein abundance differences at this age, resulting in 170 dysregulated proteins (11 up, 159 down, **fig. 4B**, **table S9**), which were largely distinct from differentially expressed proteins in TG-CTL synaptosomes at 4 months (**fig. 4C**). Upregulated proteins in 10 moa TG-CTL mice were overrepresented for 52 clusters of biological processes and 13 clusters of cellular components GO terms, mostly driven by APP-associated pathways and Aβ-associated neuroinflammation (table S10). Downregulated proteins were significantly associated with 23 clusters of BP and 17 clusters of CC, including cell communication, neuronal projection development, transport vesicle membranes, and DNA damage (**fig. S4A-B**, **table S10**).

**Figure 4.**
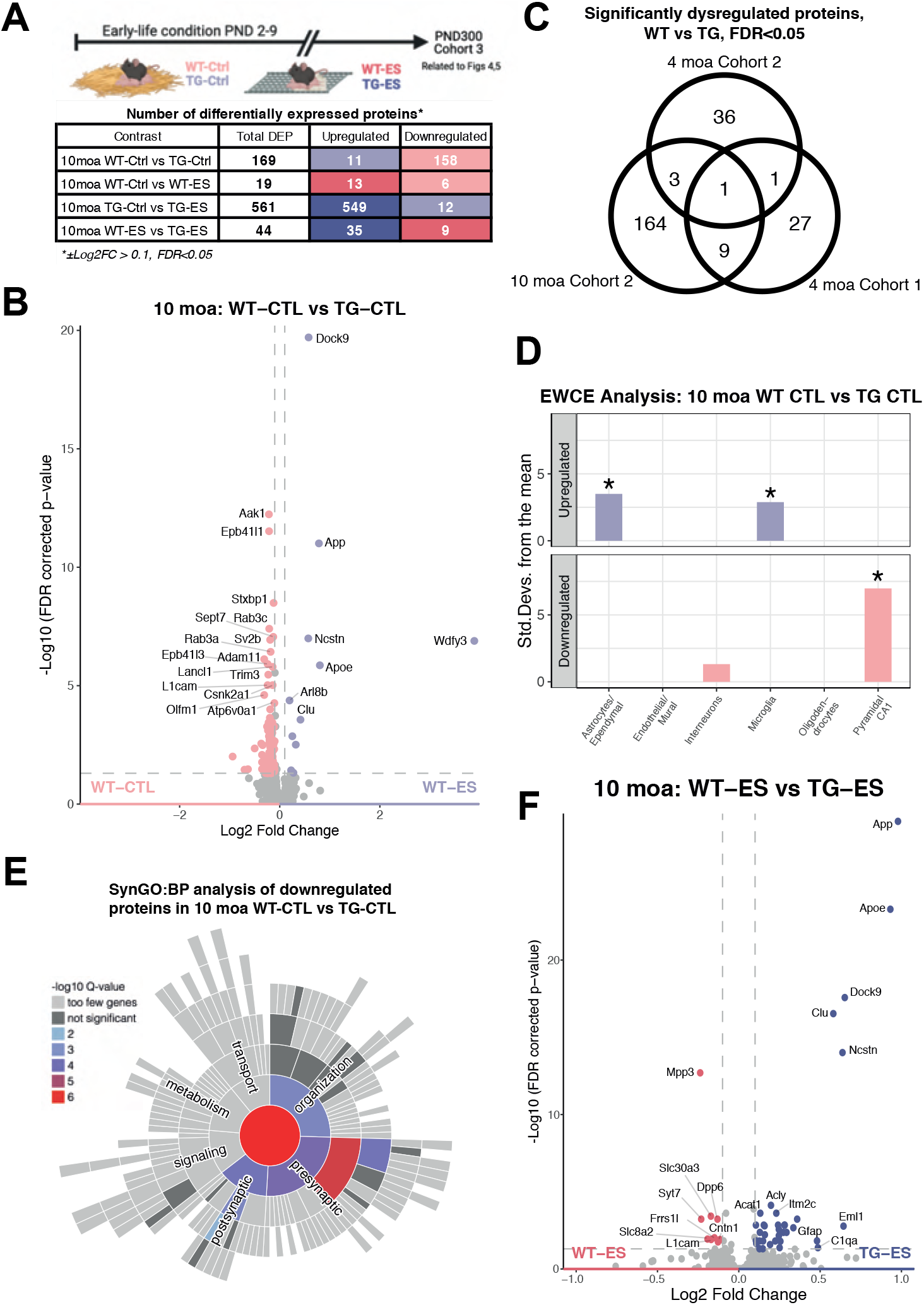
Downregulation of proteins involved synaptic transmission in hippocampal synaptosomes from APP/PS1 mice at 10 moa. (A) Experimental design within cohort 3, consisting of wildtype (WT) and transgenic APP/PS1 (TG) mice exposed to control (CTL) or early-life stress (ES) at 10 months of age (10 moa), and a summary of differentially expressed proteins across contrasts. (B) Volcano plot showing differentially expressed proteins (±Log2FC > 0.1, FDR> 0.05) in synaptosomes from TG-CTL mice at 10 moa. (C) Protein alterations in synaptosomes from 10 moa TG mice are distinct from changes in TG mice at 4 moa. (D) Expression weighted cell type enrichment (EWCE) analysis shows enrichment of astrocytic and microglial annotations in upregulated proteins in synaptosomes from TG-CTL mice. Downregulated proteins are enriched for annotations of excitatory neurons. (E) Further analysis of overrepresented biological processes in downregulated proteins using SynGO (SynGO:BP) reveal disruptions to synapse organization, presynaptic function, and postsynaptic processes. (F) Volcano plot showing differentially expressed proteins (±Log2FC > 0.1, FDR> 0.05) between in ES-exposed WT and TG mice at 10 moa.

Cell type enrichment analysis using EWCE revealed upregulated proteins to be microglial and astrocytic, whereas downregulated proteins were mostly neuronal (**fig. 4D**), consistent with the well-documented neuroinflammatory activation (68) and synaptic dysfunction (69) that results from Aβ pathology. To further investigate the synaptic pathways associated with the downregulated proteins, we performed an overrepresentation analysis for biological processes using SynGO (**fig. 4E, S4C, table S11**). This analysis revealed an under-representation of proteins in the synapses of 10 moa TG mice involved in both presynaptic vesicle exocytosis (via Git1, Ppfia3, Rab3a, Rimbp2, Rph3a, Stx1a, Stxbp1, Sv2b, Syp, Syt7, Vamp2), as well as postsynaptic organization (via Actb, Dlg1, Dlg3, Git1, Mpp2, Rimbp2) and receptor endocytosis (via Akap5, Ap2b1, Ap2m1, Ap2s1, Hpca, Synj1).

The effects of Aβ-overexpression on synaptosomal proteins were also evident when comparing ES-exposed WT and TG mice (**fig. 4F**), in which 44 proteins were differentially expressed (35 up, 9 down**, table S13**). Upregulated proteins were similarly associated with known Aβ-processing pathways, such as well as annotations to cellular components such as high-density lipoproteins, lysosomes, and mitochondria (**table S10**). Checking SynGO annotations on the 9 downregulated proteins revealed alterations to both pre-and post-synaptic compartments, with no further significant sub-terms (**fig. S4D**). Notably, 11 out of 44 differentially expressed proteins (upregulated: Apoe, App, Clu, Dock9, Gfap, Nctsn; downregulated: Lgi1, L1cam, Mpp3, Slc30a3, Syt7) were similarly differentially expressed between synaptosomes from TG-CTL and WT-CTL mice, suggesting these to be ‘core’ alterations that Aβ pathology induces on hippocampal synaptosomes at this age.

### Prior ES exposure strongly alters the proteome of hippocampal synaptosomes from APP/PS1, but not WT mice at 10 moa

We previously hypothesized that ES-associated effects at later ages would be minimal at baseline, but would be more pronounced upon exposure to subsequent challenges later in life (70). In line with this, and in contrast to the data at 4 moa, only 19 proteins (13 up, 6 down) were differentially expressed in 10 moa ES-exposed WT synapses (**fig. 5A, table S12**). Upregulated proteins did not fit into a canonically brain-related pathway, with one overrepresented biological process involving blood pressure regulation (**fig. S4E-F**). Downregulated proteins were associated with 19 clusters of GO terms (13BP, 6 CC) involving endoplasmic reticulum calcium transport (**fig. S4 E-F,** ATP2A2, RYR2, **table S10**). Also, in contrast to the data at 4 moa, where we found 18 shared differentially expressed proteins between ES and TG effects, only one protein (KCNQ2) was significantly altered in both contrasts at this age.

**Figure 5.**
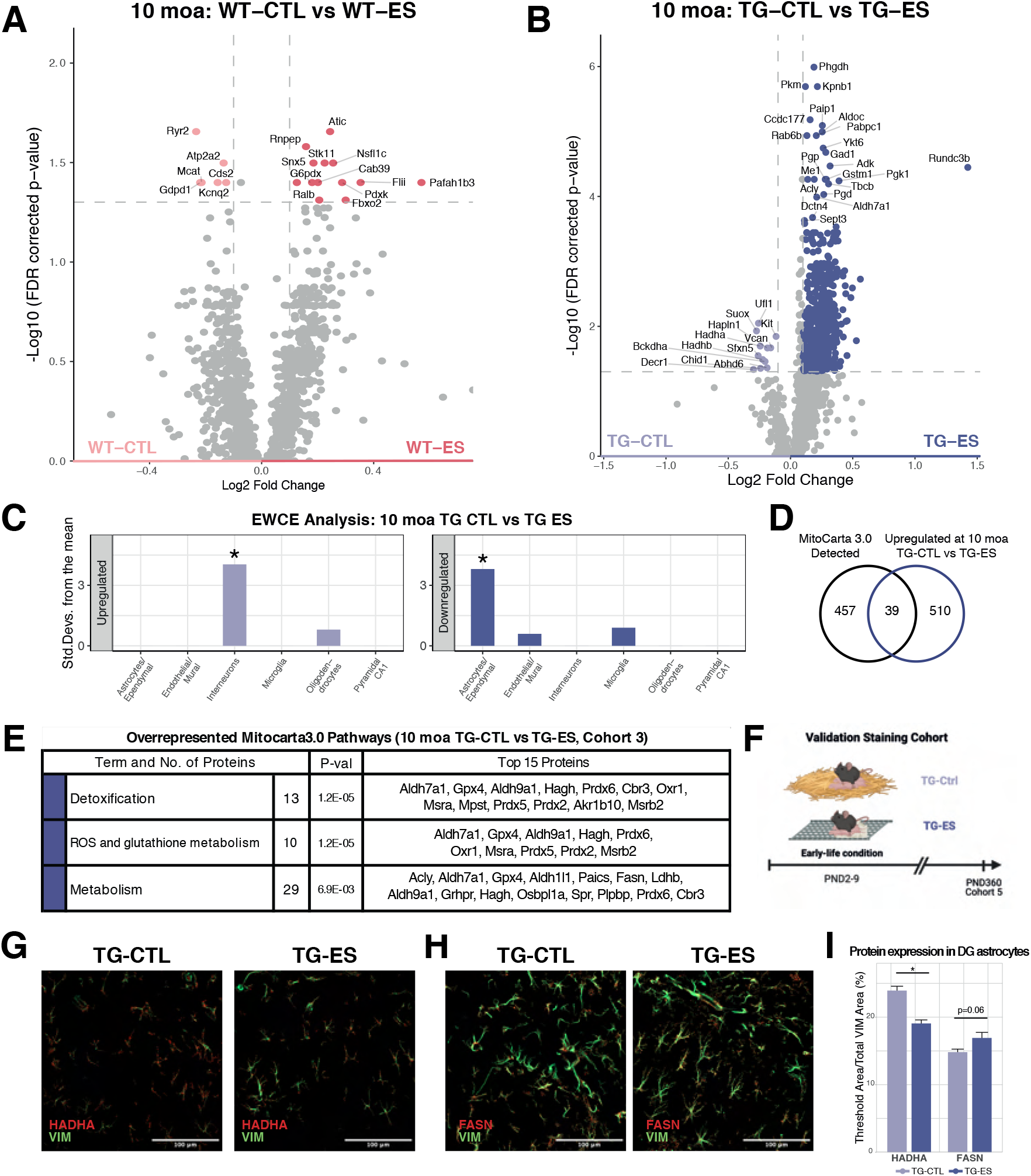
ES exposure further alters (astrocytic) mitochondrial lipid metabolism in TG synaptosomes with advanced pathology. (A) Volcano plot showing differentially expressed proteins (±Log2FC > 0.1, FDR> 0.05) in ES-exposed WT at 10 moa. (B) Volcano plot showing further differential expression in synapses from ES-exposed TG mice at this age. (C) Expression weighted cell type enrichment (EWCE) analysis shows enrichment for interneuronal annotations in upregulated proteins in ES-exposed TG mice. Downregulated proteins are enriched for astrocytic annotations. (D-E) Differentially expressed proteins in this contrast overlap with MitoCarta (D) and are functionally annotated to detoxification and reactive oxygen species metabolism (E). (F-I) Overview of cohort 5 (F), used to confirm lasting alterations to astrocytic expression of Hadha (G) and Fasn (H) within ES-exposed TG hippocampus at 12 months (I). *, condition effect, p>0.05. Analyses performed using student’s T test, Hadha – TG-CTL: n=7 TG-ES: n=5, Fasn – TG-CTL: n=8 TG-ES: n=6.

Furthermore, while ES did not have lasting effects in synaptosomes from WT mice, we found that ES-exposure massively shifted the proteomic profile of synaptosomes from TG mice at 10 moa, with 561 differentially expressed proteins (549 up, 12 down, **fig. 5B, table S14**). GO analysis of the upregulated proteins (**fig. S4G-H, table S10**) revealed terms related to the metabolism of different molecules such as carbohydrates, aldehydes, NAD, organonitrogens, organophosphates, hydroxy-compounds, as well as GO terms related to DNA metabolic processes and chromosome structure. Cellular component analysis indicated 8 clusters of GO terms that suggest alterations at the nucleus, cytosol, (microtubule) cytoskeleton, proteasome complex, and V-type proton ATPases. GO terms that were associated with downregulated proteins in CTL vs ES TG mice (**fig. S4G-H, table S10**) were involved in fatty acid metabolism and beta-oxidation in the mitochondria, driven by four proteins (Bckdha, Decr1, Hadha, Hadhb).

EWCE analysis of the differentially expressed proteins revealed the upregulated proteins to be related to interneurons and downregulated proteins to be astrocytic (**fig. 5C**). Upregulated proteins were not enriched for any SynGO annotated pathways (not shown). Additionally, ES exposure strongly affected mitochondrial proteins in TG synaptosomes at this age, with 47 out of 565 differentially expressed proteins (41 up, 6 down) overlapping with MitoCarta (**fig. 5D**, in contrast with 2 mitochondrial proteins, HMCL and MPST, altered between WT-CTL and TG-CTL synaptosomes). GO analysis on these 47 proteins using MitoCarta-annotated pathways revealed an overrepresentation of proteins associated with type II fatty acid synthesis, detoxification, ROS/glutathione metabolism and sulfur metabolism (**fig. 5E**). In conjunction with the gene ontology results, these data implicate mitochondrial alterations to underlie the effects of ES exposure on the synaptosomes of TG mice.

To validate the results from the proteomic analysis, and also investigate the temporal persistence of the presumed ES effects on lipid metabolism in the hippocampi of TG mice, we stained for Hadha and Fasn (down- and upregulated in the proteomics data, respectively) in a cohort of 12-month-old (12 moa) TG mice exposed to ES (Cohort 5, **fig. 5F**). Because of the strong expression of these two proteins in astrocytes (71,72), and the EWCE data suggesting further astrocytic dysfunction in ES-exposed TG mice, we quantified the expression of these proteins together with Vimentin as an astrocyte-specific marker (**fig. 5G, 5H**). We found a decrease in astrocytic Hadha (t(9.98)=2.2486, p=0.0483, **fig. 5I**) and a trend for increased astrocytic Fasn (t(6.422)=-2.2454, p=0.0630, **fig. 5I**), specifically within the dentate gyrus (DG) of the hippocampus. Beyond corroborating the protein data, these results also suggest that alterations to hippocampal astrocytic lipid metabolism in ES-exposed TG mice occur and may also persist beyond just the synapse.

### ES modulates the temporal pattern of synaptic mitochondrial protein alterations seen in TG mice

Lastly, we aimed to contextualize the alterations to (mitochondrial) lipid metabolism proteins in ES-exposed TG mice, by juxtaposing them against the TG-associated differential expression of these proteins progress over time. To do so, we selected MitoCarta annotated proteins that were significantly altered (±Log2FC > 0.1, FDR <0.05) in the WT-CTL vs TG-CTL and WT-CTL vs WT-ES contrasts at either age (**fig. 6A**). Doing so revealed 55 unique MitoCarta proteins, of which 48 were present in all datasets.

**Figure 6.**
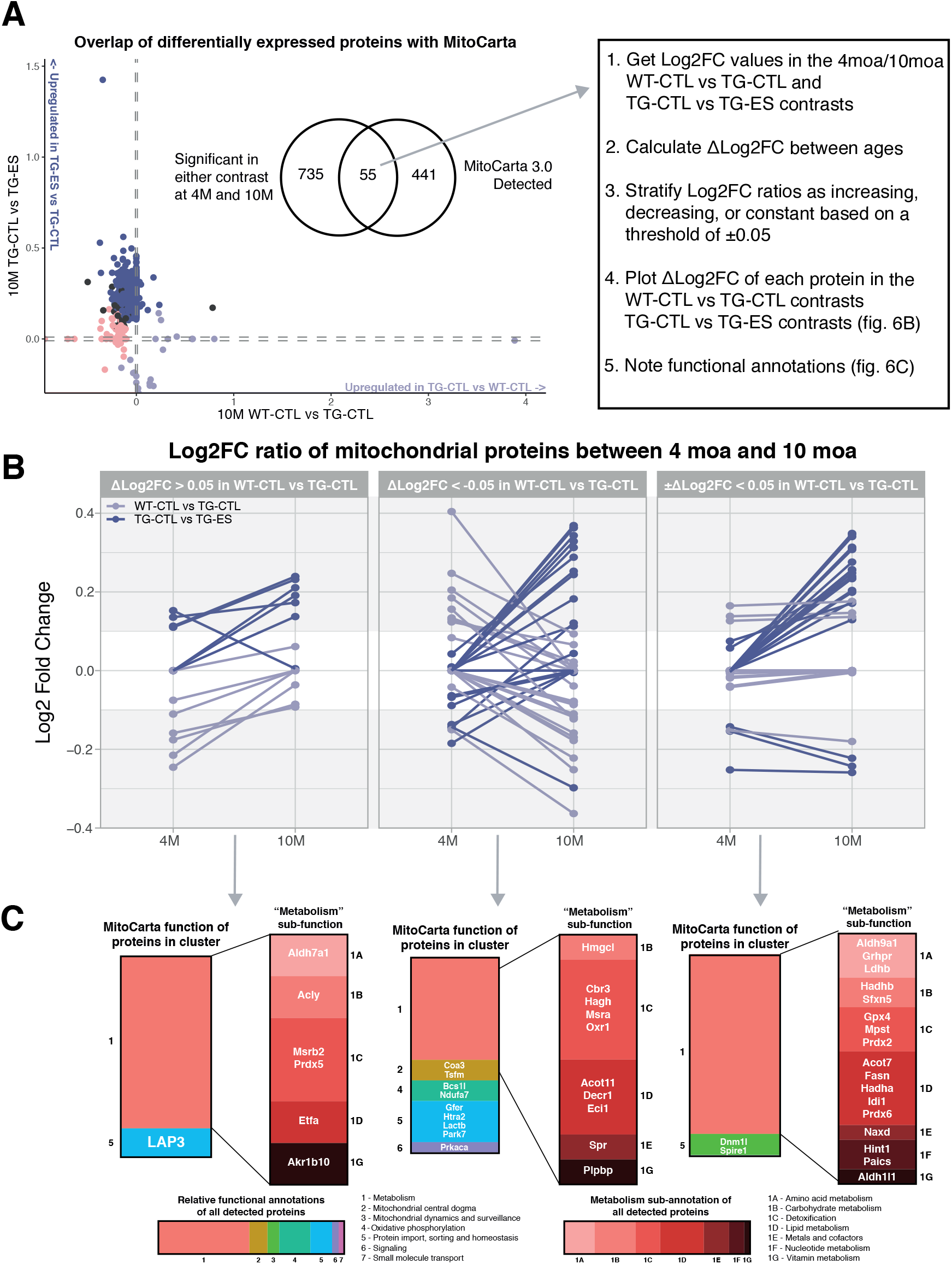
Exploring how ES exposure influences the temporal pattern of mitochondrial protein expression in synaptosomes from TG mice. (A) Workflow to visualize trajectory of mitochondrial protein alterations in TG synapses. We identified proteins significantly altered (±Log2FC > 0.1, FDR > 0.05) in synaptosomes from TG-CTL and ES-exposed TG mice from cohorts 2 and 3. A Log2FC ratio between both ages was calculated (Log2FC10moa – Log2FC4moa) to visualize the ‘standard’ progression of these changes in TG-CTL mice. (B) Based on a threshold of ΔLog2FC > ±0.05, proteins were stratified based on whether their alterations in TG mice (light blue lines) get stronger (ΔLog2FC >0.05), weaker (ΔLog2FC >-0.05), or are consistent (ΔLog2FC > ±0.05) with age. The ΔLog2FC of the same proteins in ES-exposed TG mice are then visualized (dark blue lines). (C) Functional annotations of proteins within each ‘cluster’ using MitoCarta reveal that mitochondrial proteins lastingly altered in the synaptosomes of both CTL and ES-exposed TG mice have metabolic functions, e.g. in lipid metabolism and detoxification.

These proteins were affected in different ways: for instance, of the proteins upregulated (Log2FC > 0.1) in TG mice at 4 moa, some are downregulated (Log2FC < -0.1) in the 10 moa animals (Prkaca), while some are upregulated to a similar extent (Hadha, Hadhb, Sfnx5, **table S15**). On the other hand, certain proteins downregulated in synaptosomes from 4 moa TG mice are no longer altered at 10 moa (Akr1b10, Etfa, Lap3, Msrb2, Prdx5). We thus set out to quantify the extent to which these 48 proteins are differentially expressed in synapses in TG mice with different age.

To visualize this, we calculated a ratio of the Log2 fold change in the expression of each protein when comparing synaptosomes from WT-CTL versus TG-CTL mice at 4 moa and 10 moa (ΔLog2FC = Log2FC_10moa_ – Log2FC_4moa_, adapted from (24)). This proxy measure is not able to account for the absolute magnitude of change in each protein across ages. However, it is nonetheless able to give an impression of the relative TG effects on synaptic protein expression and whether upregulation/downregulation in synapses from 4 moa TG mice remain or are further upregulated/downregulated at 10 moa. As such, proteins were stratified (using a ΔLog2FC threshold of ±0.05) by whether either the ratio of their relative expression increases (ΔLog2FC > 0.05), decreases (ΔLog2FC < -0.05), or does not change (±ΔLog2FC < 0.05) between both ages in the TG mice **(fig. 6B, light blue lines).**

We then visualized the ΔLog2FC ratio of these same proteins in the hippocampal synaptosomes, based on their Log2FC in ES-exposed TG mice **(fig. 6B, dark blue lines)**. Analyzing the functional annotations of these proteins based on MitoCarta reveal that mitochondrial aberrations in the synapses of TG mice are specifically involved in metabolism (**fig. 6C**). Notably, ES exposure seems to lead to a different temporal pattern in the expression of these proteins with proteins with a negative ΔLog2FC, as well as those of which the Log2FC ratio is not changed in TG-CTL mice. These proteins are particularly involved in lipid metabolism and (ROS) detoxification, suggesting that further ES alterations to TG synapses might act through these pathways.

## Discussion

We here studied the hippocampal synaptosomal proteome in transgenic amyloid-beta (Aβ) overexpressing APP/PS1 mice after postnatal early-life stress (ES) exposure. Interactions between ES and transgenic Aβ overexpression were studied at 4 and 10 months of age, representing early and advanced pathological stages, respectively. At 4 moa, synaptosomes isolated from both control (CTL) APP/PS1 and ES-exposed wildtype (WT) mice exhibited a similar upregulation of mitochondrial proteins and a downregulation of actin-related proteins, while ES-exposure did not further alter the synaptosomes from APP/PS1 mice. At 10 moa, proteins involved in Aβ processing were found to be upregulated, while specific neuronal proteins involved in both pre- and post-synaptic processes were downregulated. At 10 moa, while there were minimal ES effects on proteins of the synaptosomes in WT mice, ES-exposed APP/PS1 mice exhibited further alterations in the expression of synaptosomal proteins involved in lipid metabolism. Our temporal analysis on the effects of ES on the synaptic proteome indicated that aspects of the mitochondrial metabolism are lastingly altered in synaptosomes of APP/PS1, both during early pathological stages and exacerbated during more advanced pathological stages.

### Synaptic mitochondria are altered by both early pathological changes induced by transgenic overexpression of Aβ as well as by ES treatment/exposure

We found both mitochondrial proteins and morphology altered in APP/PS1 mice at an age representing an early stage of Aβ pathology. Namely, synaptosomes from APP/PS1 mice across both cohorts displayed an upregulation of mitochondrial proteins, which was accompanied by a decrease in the number of presynaptic mitochondria, as well as by smaller postsynaptic mitochondria.

Our findings support reports that mitochondrial dysfunction contributes to early Alzheimer’s disease (AD) pathogenesis (73). This is thought to be in part caused by the interactions between Aβ species and mitochondria (74), as Aβ has been shown to alter mitochondrial morphology (75,76), fusion/fission balance (77), and proteostasis (75,78). In fact, AD patients exhibit region-specific mitochondrial loss in the central nervous system (75,79,80), leading to the notion that mitochondrial alterations may drive AD pathogenesis (73,81,82).

A loss of mitochondria has major implications for synapses, which have high energy demands (83–85). A disrupted energy supply is particularly evident in the hippocampi of AD patients (86) and might be due to deficient oxidative phosphorylation and electron transport, as also found in studies using whole hippocampi of 3- and 6-month-old APP/PS1 mice (87,88). Our data suggest that these alterations occur also specifically in the synapses of APP/PS1 mice, in line with a study that found reduced numbers of mitochondria in the same model at 6 months of age (89). Importantly, the fewer pre-synaptic mitochondria we found were not larger in area, suggesting that the reduction in number is not due to increased fusion events (90), but likely due to the loss of mitochondria. As such, the increased expression of mitochondrial proteins we observed in synapses from APP/PS1 mice, especially at 4 moa, might reflect compensatory mechanisms to overcome these putative deficits. The functional implications of these changes for synaptic energy homeostasis (e.g., ATP production) remain to be determined.

Our data also highlight how ES may lead to functional mitochondrial alterations at the synapse. As in APP/PS1 mice, synaptosomes from 4 moa WT mice exposed to ES were also enriched for certain mitochondrial proteins. These changes were accompanied by a decrease in mitochondrial counts at the ultrastructural level (with no increase in mitochondrial area), which implies the loss of mitochondria. Notably, this decrease in mitochondrial count was found in the post-synapse, in contrast to the pre-synaptic mitochondrial deficits in APP/PS1 synapses. The upregulation in mitochondrial proteins might then, analogously, be compensatory against the effects of such deficits. This would explain a previous finding of increased fatty acid synthase (Fasn) mRNA expression in the whole lysates from the contralateral hippocampus of the same WT-ES mice used in this study (30). This link between stress and mitochondrial function is in line with a study showing the impairment of respiratory processes in synaptosomal mitochondria of mice subjected to chronic stress in adulthood (91). Additionally, we have previously reported an ES-mediated disruption of the electron transport chain activity in the hypothalamus and muscle tissues in postnatal pups, which persisted until at least 10 months of age (92). That same study reported an age-associated change in expression of hippocampal mitochondrial fission proteins after ES, with Fis1 being less expressed in ES-exposed pups, yet more highly expressed in 10 moa WT-ES mice.

Crucially, we also found evidence for an association between the effects of ES exposure and APP/PS1 genotype on hippocampal synaptic mitochondria. As discussed above, the effects of ES on synaptic mitochondria seem to mirror the changes occurring in APP/PS1 synapses. Importantly, the degree to which they overlap and converge needs further investigation, as does the extent of mitochondrial involvement in how ES effects impacts APP/PS1 mice. The interaction between the two experimental factors is particularly pronounced at 10 moa, where 39 mitochondrial proteins were upregulated in ES-exposed APP/PS1 mice (compared to only 2 when comparing WT and APP/PS1 mice at this age). These mitochondrial disruptions might possibly be caused by the increased Aβ plaque load observed in ES mice at this age (29), as amyloid plaques have been shown to be at least focal sources of toxicity that impair mitochondrial structure and function (93).

### Actin dynamics as a substrate for synaptic alterations in both APP/PS1 and ES-exposed WT mice

We also found evidence for a downregulation of actin-dynamics-related processes in APP/PS1 mice at 4 moa. This seemed to be mediated by the SCAR Complex, an evolutionarily conserved family of proteins involved in de novo assembly of actin branches (56,94). Furthermore, downregulated proteins in APP/PS1 synapses are also involved in the regulation of actin polymerization and depolymerization.

The actin cytoskeleton is important for synaptic structure, being involved e.g., in the formation, maintenance, or elimination of dendritic spines, as it dynamically reorganizes in response to synaptic signals (95). Importantly, there is evidence for dysfunctional actin in the brains of AD patients and mouse models, seen as neuropathological ‘rods’ formed by actin-depolymerizing factor (ADF) and Cofilin ‘rod’ aggregates (96). These aggregates can be induced via cellular exposure to stressors such as reactive oxygen species (ROS) and Aβ, and form along mitochondria-deficient parts of the neurite (97,98). This process, involving Cofilin-mediated formation of ‘rods’ and arrests to actin (de)polymerization, is thought to be part of the cellular response to stress (99), reducing the rate of ATP-loss (100), which might be explained by the high energetic costs associated with actin disassembly (101).

Mitochondrial trafficking in part also occurs via actin networks (102), and mitochondrial functions (e.g., fusion/fission) are also influenced by actin remodeling (103). As such, it is tempting to speculate that the potential alterations to actin dynamics at the synapse is a result of a diversion of energy expenditure to meet energetic demands brought about by e.g. a loss of mitochondria at this age. Although this might have energetic benefits on the short term, it is important to consider that it might lead to long term disruptions. While Aβ-induced alterations in actin-related protein levels are most prominent in synaptosomes from 4 moa APP/PS1 mice, changes in these are still apparent at 10 moa (**table S3 and S10**). As actin dynamics are also important for the trafficking of receptors to the synapse (104,105), as well as synaptic vesicle release and recycling (106,107), we speculate that the prolonged effect on proteins relating to these processes might possibly explain the downregulation of presynaptic release and postsynaptic receptor endocytosis in synapses from 10 moa APP/PS1 mice.

It is intriguing to further consider how ES might impact cytoskeletal dynamics, given that it was also affected in the synaptosomes isolated from ES-exposed WT mice at 4 moa. The potential effects on the actin cytoskeleton are in line with studies on early life social isolation in rats. Here, it was shown that a decrease in actin turnover by ADF/cofilin inactivation, led to an impaired trafficking of AMPA receptors (108,109). These mechanisms might explain the altered distribution of receptors that have been described in the synapses of mice exposed to ES using the same limited nesting model we used (33). Moreover, disruption of actin dynamics will have consequences beyond the neural compartments of the synapse, e.g., in the motility of perisynaptic astrocytes (PAP), which are known to undergo cytoskeletal remodeling in response to neuronal activity (45,110). Crucially, PAP motility decreases as a function of astrocytic coverage around dendritic spines (111), in line with our data showing an upregulation of astrocyte-associated proteins in synaptosomes from WT-ES mice. The presence of less motile PAPs in ES synapses would be expected to have consequences in their ability to e.g. regulate extracellular neurotransmitter uptake and gliotransmitter release (110,112), and thereby overall synaptic strength and health. While these might be a consequence of the deficits in synaptic structure (113–116) and plasticity (33,117) described in various ES models, their possible role in exacerbating synaptic impairments in ES synapses remain to be seen.

Separately, while the type of downregulated actin-related proteins in ES mice overlapped considerable with those from APP/PS1 mice, the downregulated proteins in ES mice were flagged by EWCE to be enriched in oligodendrocytic annotations, which is caused by proteins that have also been detected in other synaptosomal preparations (24,45,110,118). Actin dynamics also plays an important role in the formation of growth cones that are important for the maturation and migration of cells of an oligodendrocyte lineage (119), which have been reported to form synaptic contacts with both excitatory (120) and inhibitory (121) neurons in the hippocampus. Cells from this lineage have also been reported to be affected by ES exposure (122,123), and potentially represent another substrate through which actin dynamics are altered in ES mice. It is currently not clear whether proteins originating in these cells can end up in a synaptosomal preparation. That said, our data altogether suggest a contribution of cytoskeletal alterations in APP/PS1 and ES phenotypes. The extent to which these changes relate to those found in mitochondria, overlap between these experimental treatments, and are viable targets for preventive/intervention strategies, remains to be investigated in the future.

### ES effects on the trajectory of Aβ-induced synapse pathology: impact on (astrocytic) mitochondrial metabolism

While ES exposure did not alter protein expression in synapses from APP/PS1 mice at 4 moa, it did lead to vast differences in protein expression at 10 moa. Because we studied mice from these experimental groups at both 4 moa and 10 moa, we also explored how ES impacted the trajectory of Aβ-associated changes in hippocampal synaptosomes. We focused our current analysis on mitochondria, given its prominence in the proteomic data, aiming to provide evidence that ES exposure modulates the TG effects on these proteins.

The idea that ES could modulate AD- and other aging-associated trajectories is not new (70). We have previously shown that ES exposure has age-dependent effects on Aβ pathology in AD mice, decreasing cell-associated amyloid at 4 moa, and increasing Aβ plaque load in the hippocampus of stressed TG mice at 10 moa (29). Similarly, ES exposure increases the expression of amyloidogenic proteins such as BACE1 in the hippocampus of 6- and 12-month-old TG mice (31). This has also been studied in the context of microglia and astrocytes, where we found, e.g., that ES leads to age-dependent alterations in microglial morphology and gene expression in APP/PS1 mice (29,30). The synaptic alterations we detected, especially in ES-exposed APP/PS1 mice, likely reflect the age-associated increase in Aβ pathology and the indirect effects that occur as a result of the continuous exposure to Aβ.

We hypothesized that the lack of additive ES effects in 4 moa APP/PS1 mice is indicative of a convergence between ES effects with genotype effects at this age. The lack of differentially expressed proteins is consistent with reported effects of stress exposure during different life stages on AD-like pathology (124). For instance, maternally separated WT rats show upregulated protein expression of BACE1 and Aβ (6), and mice exposed to chronic psychosocial stress similarly increased expression of AD-associated proteins (4). We partly explored this hypothesis by EM analysis of mitochondria in both ES-exposed WT and APP/PS1 mice, where we found decreased numbers of presynaptic mitochondria in APP/PS1 hippocampi, and fewer mitochondria in the post-synapse of WT-ES mice. The mitochondrial dysfunction at these respective synaptic compartments might result in energetic consequences that are not further impaired in ES-exposed APP/PS1 mice. This will need to be functionally demonstrated in further studies.

In addition, ES exposure strongly affected synaptosomes from APP/PS1 mice at 10 moa, when Aβ is more severe and widespread (125). This is in contrast to minimal changes in protein expression in synaptosomes from 10 moa WT mice, supporting the hypothesis that ES effects at later ages would necessitate secondary challenges (as Aβ would represent) to be unmasked (70). These effects were prominent in the dysregulation of mitochondrial proteins, which were involved in ROS as well as fatty acid metabolism. Strikingly, these processes seem to be particularly affected in astrocytes, in line with the prominent role of these cell types in energy balance in the brain (72). Astrocytes were also indicated to be affected in our EWCE analysis of differentially expressed proteins across several contrasts (**fig. 2C, 4C, 5C, S1A**), in line with astrocytic protein detection in other synaptosomal studies (24,45,110,118).

Finally, our data highlight the importance of astrocytes in mediating both APP/PS1 and ES effects (30,126). We have previously speculated that astrocytes in ES-exposed APP/PS1 mice, while similarly reactive to control APP-PS1 as quantified by GFAP immunostaining, might also exhibit alterations in other functional domains (e.g. lipid metabolism, blood-brain barrier maintenance), due to the increased plaque load in their surroundings (30). Here, we show evidence that astrocytic lipid metabolism is further disrupted in ES-exposed APP/PS1 mice. For instance, we find evidence for decreased astrocytic expression of Hadha, a key enzyme in the beta-oxidation of fatty acids (72), as well as increased astrocytic expression of Fasn, an enzyme involved in de novo synthesis of fatty acids (71). These changes seemed to occur both at the synapse at 10 moa and in the rest of the dentate gyrus, as validated with immunofluorescence at 12 months, possibly indicating a shift in lipid homeostasis. Beyond indicative of a compensatory response to energy deficits, altered lipid metabolism in the hippocampi of ES-exposed APP/PS1 mice could have consequences for neuronal circuits, given the recent evidence that the contents of saturated lipids from reactive astrocytes can have neurotoxic properties (127). At the same time, an imbalance in the synthesis of fatty acids may mean increased rates of oxidative stress, given their propensity to be peroxidized by free radicals (128). This seemed to occur predominantly in astrocytes, in line with the important role of their mitochondria in energy production (72,129).

In this context, our observation that mitochondrial proteins associated with ROS metabolism are upregulated in the synapses of ES-exposed APP/PS1 mice, might be a compensatory adaptation brought about by this shift in lipid synthesis. A similar effect was found in work using the 5xFAD TG model (130), suggesting that this is induced by Aβ pathology. Importantly, we have previously shown age-dependent consequences of ES on fatty acid profiles across brain regions in WT mice (131,132), where e.g. hippocampal polyunsaturated fatty acids (PUFA) are decreased in ES-exposed pups and increased in ES-exposed adult mice at 6 months (132). Notably, the alteration of PUFA species, via dietary intervention, has been shown to rescue both lipid levels and cognitive deficits in ES-exposed adults (132). The consequences of these altered lipid profiles in the ES-exposed brain (e.g., regarding ROS metabolism), as well as the functional consequences of ES on these processes in the APP/PS1 brain, will require further investigation.

## Conclusion

Our work revealed that ES modulates Aβ pathology-induced alterations in the synaptosomal proteome. We show that the effects of ES alone on the synaptic proteome are similar as those of early Aβ pathology, and accordingly, ES exposure did not additionally alter the proteomic profile in APP/PS1 mice at this age. We also report that ES already leads to additional effects on the proteomic alterations, similar to those caused by Aβ at advanced pathological stages, which was minimal in the WT condition. These data highlight the importance of the synapse as a long-term substrate for APP/PS1 pathology and show how ES influences the trajectory of these alterations with advancing age. These interactions between the two factors also suggest that Aβ pathology, especially early on, leads to a synaptic signature similar to that after early stress exposure.

## Declarations

### Ethical Approval and Consent to participate

Not applicable

### Consent for publication

All co-authors have read the final version of the manuscript, and consent to its submission for publication.

### Availability of supporting data

Our proteomics data are publicly available and can be accessed online using a web app, via https://amsterdamstudygroup.shinyapps.io/ES_Synaptosome_Proteomics_Visualizer/. Tables containing gene lists and results of (downstream) proteomics analyses are included as supplementary files. All other data will be made available upon request.

## Author contributions

- Study conceptualization: AK, MHGV, JMK, MSJK
- Animal work and tissue collection: JMK, MSJK, LH, NB, SLL
- Sample preparation and mass spectrometry: MSJK
- Downstream analysis of mass spectrometry data: JMK, MSJK
- EM sample preparation and imaging: MSJK
- EM analysis: MSJK and RT
- Validation staining, imaging, analysis: LM, EY
- Figures and tables: JMK
- Manuscript drafting and writing: JMK, AK
- Manuscript review and editing: JMK, MSJK, SLL, PJL, ABS, HK, AK, MHGV
- All authors have read the final version of the manuscript.

## Acknowledgments

Not applicable

## Authors’ information

Not applicable

## Abbreviations

ES: early-life stress
AD: Alzheimer’s disease
WT: wildtype CTL: control
4 moa: 4 months of age
10 moa: 10 months of age
TG: transgenic APP/PS1 genotype
Aβ: amyloid-β

---

**Figure S1. Confirmation of synaptic proteins in synaptosomal isolation**

(A) Proteins detected across the three cohorts of proteomics experiments highly overlapped and were also mapped to a majority of the 1113 proteins annotated in SynGO v1.1. (B) Sunburst plots of cellular components (CC, top row) and biological processes (BP, bottom row) enriched in SynGO-annotated detected proteins across three cohorts compared to a background of brain expressed genes as defined by SynGO. (C) Top 15 of the overrepresented CC and BP terms (as arranged by p-values) when performing an overrepresentation of all detected proteins against the same background of brain expressed genes using gProfiler2.

**Figure S2. Overview of (top) GO pathways altered by APP/PS1 genotype and ES exposure at 4 moa (related to figs. 1, 2)**

(A) Expression weighted cell type enrichment (EWCE) analysis in Cohort 1 shows upregulation of astrocytic proteins and downregulation of neuronal proteins TG vs WT synaptosomes. (B) EWCE analysis in WT-CTL vs TG-CTL synaptosomes from Cohort 2 reveal downregulation of neuronal proteins. (C) Sunburst plot showing significant SynGO-annotated biological processes (SynGO:BP) in downregulated proteins between the WT-CTL vs TG-CTL contrast in Cohort 2. (D) Table of overrepresented SynGO:BP terms depicted in panel C, as well as a table of overrepresented SynGO-annotated cellular component (SynGO:CC) terms. (E-F) Top GO terms overrepresented in significantly upregulated (E) and downregulated (F) proteins in 4 moa WT-CTL vs WT-ES synaptosomes. (G) Overrepresented SynGO:BP terms in downregulated proteins in synaptosomes from WT-ES vs TG-ES mice at 4 moa. All GO analyses performed based on biological processes (BP) and cellular components (CC). Italicized terms are ‘parent’ terms after clustering by semantic similarity using RRVGO.

**Figure S3. Other measures of mitochondrial morphology are unaffected in ES or TG synapses at 4 moa (related to fig. 3)**

(A) Total number of mitochondria is not affected in the hippocampi of ES-exposed WT and TG mice. (B) Mitochondrial area in the dendritic (DEND), and astrocytic (ASTRO) compartments are not affected at 4 moa. (C-D) Mitochondrial perimeter (C) and circularity (D) in the PRE, post-synaptic (POST), DEND, and ASTRO compartments are not altered at 4 moa.

**Figure S4. Overview of (top) overrepresented pathways altered by APP/PS1 genotype and ES exposure at 10 moa (related to figs. 4, 5).**

(A) Top overrepresented gene ontology biological processes (GO:BP) terms from upregulated (light blue) and downregulated (light red) proteins when comparing synaptosomes from 10 moa WT-CTL and TG-CTL mice. (B) Top overrepresented GO cellular component (GO:CC) terms from upregulated (light blue) and downregulated (light red) proteins when comparing synaptosomes from 10 moa WT-CTL and TG-CTL mice. (C) SynGO analysis of biological processes (SynGO:BP) overrepresented in downregulated proteins (light red) in 10 moa TG-CTL vs WT-CTL mice reveals alterations to pre-synaptic vesicle cycling and post-synaptic receptor endocytosis. (D) Sunburst diagram from SynGO:BP and cellular component (SynGO:CC) analysis of downregulated proteins in WT-ES vs TG-ES mice at 10 moa. (E-F) Top overrepresented GO:BP (E) and GO:CC (F) terms from upregulated (dark red) and downregulated (light red) synaptosomal proteins in ES-exposed WT mice. (G-H) Top overrepresented GO:BP (G) and GO:CC (H) terms from upregulated (dark blue) and downregulated (light blue) proteins in synaptosomes from ES-exposed TG mice. All GO analyses performed based on biological processes (BP) and cellular components (CC). Italicized terms are ‘parent’ terms after clustering by semantic similarity using RRVGO.

## Supplementary Tables

**Table S1** – Table of differentially expressed proteins between WT vs TG synaptosomes at 4 moa (Cohort 1)

**Table S2** – Table of differentially expressed proteins between WT-CTL vs TG-CTL synaptosomes at 4 moa (Cohort 2)

**Table S3** – Table of overrepresented gene ontology (GO) terms and reduced parent terms from differentially expressed proteins at 4 moa (Cohorts 1 and 2)

**Table S4** – Table of differentially expressed proteins between WT-CTL vs WT-ES synaptosomes at 4 moa (Cohort 2)

**Table S5** – Table of differentially expressed proteins between TG-CTL vs TG-ES synaptosomes at 4 moa (Cohort 2)

**Table S6** – Table of differentially expressed proteins between WT-ES vs TG-ES synaptosomes at 4 moa (Cohort 2)

**Table S7** – Table of shared differentially expressed proteins between the WT-CTL vs TG-CTL and WT-CTL vs WT-ES contrasts at 4 moa (Cohort 2)

**Table S8** – Table of overrepresented gene ontology (GO) terms and reduced parent terms from shared differentially expressed proteins between the WT-CTL vs TG-CTL and WT-CTL vs WT-ES contrasts at 4 moa (Cohort 2)

**Table S9** – Table of differentially expressed proteins between WT-CTL vs TG-CTL synaptosomes at 10 moa (Cohort 3)

**Table S10** – Table of overrepresented gene ontology (GO) terms and reduced parent terms from differentially expressed proteins at 10 moa (Cohort 3)

**Table S11** – Table of overrepresented SynGO-annotated ontology terms from downregulated proteins in WT-CTL vs TG-CTL synaptosomes at 10 moa (Cohort 3)

**Table S12** – Table of differentially expressed proteins between WT-CTL vs WT-ES synaptosomes at 10 moa (Cohort 3)

**Table S13** – Table of differentially expressed proteins between WT-ES vs TG-ES synaptosomes at 10 moa (Cohort 3)

**Table S14** – Table of differentially expressed proteins between TG-CTL vs TG-ES synaptosomes at 10 moa (Cohort 3)

**Table S15** – Table of Log2FC values and functional annotations of mitochondrial proteins in the WT-CTL vs TG-CTL and TG-CTL and TG-ES contrasts at 4 moa and 10 moa

