## Supplementary Figures 1-4 for "Early-life stress exposure impacts the hippocampal synaptic proteome in a mouse model of Alzheimer’s disease: age- and pathology-dependent effects on mitochondrial proteins": 20230301_Figure_S2_redo-compressed.pdf

### Suppl. Fig 2

### A

EWCE Analysis: 4 moa WT vs TG (Cohort 1)

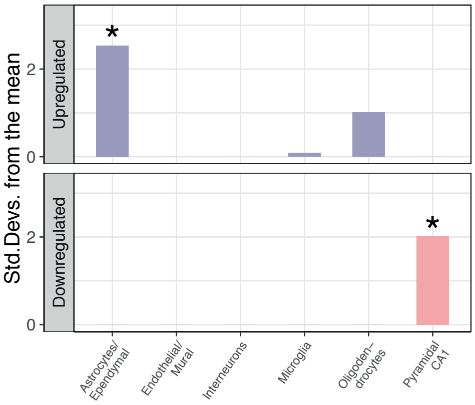


### B

EWCE Analysis: 4 moa WT-CTL vs TG-CTL (Cohort 2)

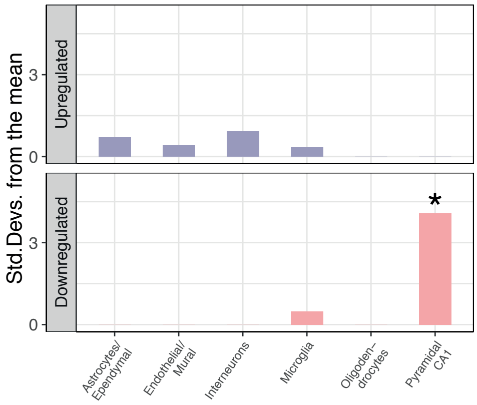


### C

SynGO:BP analysis of downregulated proteins in 4 moa WT-CTL vs TG-CTL

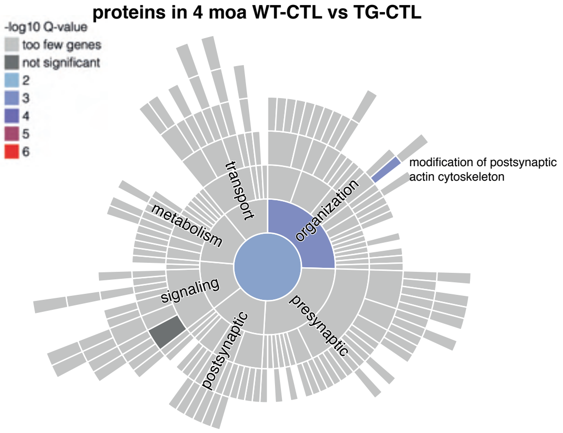


### D

**Overrepresented SynGO:BP Terms in Downregulated Proteins (4 moa WT-CTL vs TG-CTL, Cohort 2)**

| Term | Size | P-val | Proteins |
|---|---|---|---|
| process in the synapse | 10 | 3.7E-03 | Nsf, Abi1, Hpca, Pfn1, Pfn2, Ppp3ca, Wasf1, Nckap1, Add2, Dbn1 |
| chemical synaptic transmission | 3 | 3.4E-02 | Pfn1, Pfn2, Ppp3ca |
| synapse organization | 7 | 8.9E-04 | Wasf1, Pfn1, Pfn2, Nckap1, Add2, Abi1, Dbn1 |
| modification of postsynaptic actin cytoskeleton | 3 | 8.9E-04 | Wasf1, Pfn1, Pfn2 |

**Overrepresented SynGO:CC Terms in Downregulated Proteins (4 moa WT-CTL vs TG-CTL, Cohort 2)**

| Term | Size | P-val | Proteins |
|---|---|---|---|
| synapse | 12 | 2.4E-03 | Ppp3ca, Pfn2, Pfn1, Add2, Nsf, Nckap1, Wasf1, Hpca, Dbn1, Abi1, Ppp1r9b, Capzb |
| postsynapse | 10 | 7.9E-04 | Nckap1, Wasf1, Pfn2, Pfn1, Hpca, Add2, Dbn1, Abi1, Ppp1r9b, Capzb |
| postsynaptic density | 4 | 3.7E-02 | Abi1, Ppp1r9b, Dbn1, Capzb |

### E

**Top 5 Overrepresented GO:BP Terms (4 moa WT-CTL vs WT-ES, Cohort 2)**

| Term* | Size | RRVGO* | P-val | (Top 15) Proteins |
|---|---|---|---|---|
| regulation of biomineral tissue development | 11 | No | 3.5E-02 | Slc8a1, Zmpste24, Gpm6b, Gja1 |
| sodium ion homeostasis | 7 | No | 3.5E-02 | Gnai2, Atp1b2, Slc8a1, Atp1a2, Slc1a3 |
| regulation of biomineralization | 3 | No | 3.5E-02 | Slc8a1, Zmpste24, Gpm6b, Gja1 |
| regulation of bone mineralization | 5 | No | 3.5E-02 | Slc8a1, Zmpste24, Gpm6b, Gja1 |
| sodium ion export across plasma membrane | 6 | No | 3.5E-02 | Atp1b2, Slc8a1, Atp1a2, Slc4a4 |
| *negative regulation of protein polymerization* | 10 | Yes (12) | 4.4E-07 | Capza2, Capzb, Tmod2, Pfn2, Capza1, Add2, Mapre1, Cfl1, Twf2, Fkbp4 |
| *actin polymerization or depolymerization* | 12 | Yes (19) | 5.3E-07 | Ppp1r9b, Capza2, Capzb, Dbn1, Tmod2, Pfn2, Capza1, Add2, Abi2, Cfl1, Bin1, Twf2 |
| *regulation of protein polymerization* | 12 | Yes (17) | 1.0E-06 | Capza2, Capzb, Dbn1, Tmod2, Pfn2, Capza1, Add2, Mapre1, Cfl1, Bin1, Twf2, Fkbp4 |
| *regulation of actin filament polymerization* | 10 | Yes (7) | 1.5E-06 | Capza2, Capzb, Dbn1, Tmod2, Pfn2, Capza1, Add2, Cfl1, Bin1, Twf2 |
| *regulation of cellular component size* | 11 | Yes (2) | 1.5E-04 | Capza2, Capzb, Dbn1, Tmod2, Pfn2, Capza1, Add2, Cfl1, Bin1, Hsp90ab1, Twf2 |

**Italics: RRVGO analysis, parentheses indicate number of child GO terms inside cluster*

### F

**Top 5 Overrepresented GO:CC Terms (4 moa WT-CTL vs WT-ES, Cohort 2)**

| Term* | Size | RRVGO* | P-val | (Top 15) Proteins |
|---|---|---|---|---|
| *integral component of membrane* | 55 | CC (6) | 4.0E-09 | Slc4a3, Coa3, Ndufa3, Atp1b2, Cadm4, Tmem109, Lrrc8b, Cyb5a, Nlgn2, Slc3a2, Slc8a1, Stx1b, Ncam2, Ntrk2, Ptdss1 |
| *membrane* | 75 | CC (1) | 2.8E-06 | Slc4a3, Coa3, Ndufa3, Gnai2, Rab31, Snap23, Atp1b2, Lancl1, Cadm4, Tmem109, Lrrc8b, Cyb5a, Nlgn2, Slc3a2, Slc8a1 |
| *endoplasmic reticulum membrane* | 20 | CC (3) | 3.3E-04 | Tmem109, Lrrc8b, Cyb5a, Ptdss1, Lnpk, Zmpste24, Tmx1, Agpat3, Ildr2, Canx, Rpn1, Bcap31, Dnajb14, Tecr, Cyp2d22 |
| *nuclear outer membrane-endoplasmic reticulum membrane network* | 20 | CC (2) | 4.6E-04 | Tmem109, Lrrc8b, Cyb5a, Ptdss1, Lnpk, Zmpste24, Tmx1, Agpat3, Ildr2, Canx, Rpn1, Bcap31, Dnajb14, Tecr, Cyp2d22 |
| *plasma membrane* | 49 | CC (2) | 2.4E-03 | Slc4a3, Gnai2, Snap23, Atp1b2, Lancl1, Lrrc8b, Nlgn2, Slc3a2, Slc8a1, Stx1b, Ncam2, Ntrk2, Nrxn1, Grm3, Cacna2d2 |
| *mitochondria-associated endoplasmic reticulum membrane* | 3 | CC (1) | 3.3E-02 | Canx, Bcap31, Tmx2 |
| *F-actin capping protein complex* | 4 | CC (5) | 2.0E-04 | Capza2, Capzb, Capza1, Add2 |
| *actin-based cell projection* | 8 | CC (8) | 1.8E-03 | Ppp1r9b, Dbn1, Atp6v1a, Abi2, Atp6v1b2, Ca2, Cfl1, Twf2 |
| *vacuolar proton-transporting V-type ATPase, V1 domain* | 3 | CC (6) | 1.0E-02 | Atp6v1a, Atp6v1b2, Atp6v1h |
| *cortical cytoskeleton* | 6 | CC (2) | 1.2E-02 | Ppp1r9b, Capza2, Capzb, Dbn1, Mapre1, Cfl1 |
| *growth cone* | 7 | CC (3) | 2.1E-02 | Ppp1r9b, Dbn1, Tmod2, Cfl1, Hsp90ab1, Twf2, Fkbp4 |

**Italics: RRVGO analysis, parentheses indicate number of child GO terms inside cluster*

### G

**Overrepresented SynGO:BP Terms in Downregulated Proteins (WT-ES vs TG-ES, 4 moa)**

| Term and No. of Proteins | | P-val | Proteins in term |
|---|---|---|---|
| process in the postsynapse | 4 | 3.7E-03 | Hcn1, Ctnnd2, Agap3, Snap23 |
| regulation of postsynaptic membrane neurotransmitter receptor levels | 3 | 6.3E-03 | Ctnnd2, Agap3, Snap23 |
| synapse organization | 3 | 4.2E-02 | Shank2, Ctnnd2, Il1rap |
