## Supplementary Figures 1-4 for "Early-life stress exposure impacts the hippocampal synaptic proteome in a mouse model of Alzheimer’s disease: age- and pathology-dependent effects on mitochondrial proteins": 20230301_Figure_S4_redo-compressed.pdf

### Suppl. Fig 4

**A**

**Top 5 Overrepresented GO:BP Terms (10 moa WT-CTL vs TG-CTL, Cohort 3)**

| Term* | Size | RRVGO* | P-val | (Top 15) Proteins |
|---|---|---|---|---|
| regulation of tau-protein kinase activity | 3 | Yes (12) | 1.4E-04 | App, Apoe, Clu |
| protein import | 3 | Yes (22) | 1.4E-04 | App, Apoe, Clu |
| regulation of amyloid-beta clearance | 3 | Yes (3) | 2.3E-04 | App, Apoe, Clu |
| amyloid precursor protein catabolic process | 4 | Yes (8) | 4.3E-04 | App, Ncstn, Apoe, Clu |
| membrane protein ectodomain proteolysis | 3 | Yes (7) | 4.9E-04 | App, Ncstn, Apoe |
| *plasma membrane bounded cell projection morphogenesis* | 41 | Yes (11) | 5.8E-08 | Stxbp1, Rab3a, Epb41l3, L1cam, Olfm1, Chl1, Sptbn4, Akap5, Rapgef2, Map6, Actr3, Rasal1, Src, Enah, Nrcam |
| *signaling* | 90 | Yes (4) | 2.5E-07 | Aak1, Stxbp1, Sv2b, Rab3a, Csnk2a1, Olfm1, Arpc2, Lin7a, Sptbn4, Dock4, Syngr1, Ank2, Lin7c, Caskin1, Akap5 |
| *cell communication* | 90 | Yes (4) | 4.3E-07 | Aak1, Stxbp1, Sv2b, Rab3a, Csnk2a1, Olfm1, Arpc2, Lin7a, Sptbn4, Dock4, Syngr1, Ank2, Lin7c, Caskin1, Akap5 |
| *neuron projection development* | 45 | Yes (13) | 5.0E-06 | Stxbp1, Rab3a, Epb41l3, L1cam, Olfm1, Chl1, Sptbn4, Akap5, Rapgef2, Psd, Map6, Actr3, Rasal1, Strn, Enah |
| *plasma membrane bounded cell projection organization* | 50 | Yes (15) | 9.5E-06 | Stxbp1, Rab3a, Epb41l3, L1cam, Olfm1, Arpc2, Chl1, Sptbn4, Akap5, Prnp, Rapgef2, Psd, Map6, Actr3, Rasal1 |

*Italics: RRVGO analysis, parentheses indicate number of child GO terms inside cluster

**B**

**Top 5 Overrepresented GO:CC Terms (10 moa WT-CTL vs TG-CTL, Cohort 3)**

| Term* | Size | RRVGO* | P-val | (Top 15) Proteins |
|---|---|---|---|---|
| *high-density lipoprotein particle* | 3 | Yes (11) | 8.7E-06 | App, Apoe, Clu |
| *vacuole* | 7 | Yes (8) | 2.1E-05 | App, Ncstn, Wdfy3, Apoe, Arl8b, Psap, Gfap |
| *inclusion body* | 3 | Yes (2) | 2.0E-03 | Wdfy3, Clu, Psap |
| *spindle midzone* | 2 | Yes (1) | 9.7E-03 | App, Arl8b |
| *extracellular matrix* | 3 | Yes (2) | 1.3E-02 | Apoe, Clu, Vwa5a |
| *synapse* | 81 | Yes (17) | 1.7E-11 | Aak1, Stxbp1, Rab3c, Sv2b, Rab3a, Epb41l3, Trim3, Olfm1, Atp6v0a1, Arpc2, Lin7a, Syngr3, Dlg3, Syngr1, Ank2 |
| *cell projection* | 78 | Yes (9) | 4.8E-08 | Aak1, Stxbp1, Sv2b, Rab3a, Epb41l3, Trim3, L1cam, Olfm1, Arpc2, Chl1, Dlg3, Sptbn4, Dock4, Ank2, Akap5 |
| *cell periphery* | 92 | Yes (7) | 1.4E-06 | Aak1, Stxbp1, Rab3c, Rab3a, Epb41l3, Lancl1, L1cam, Csnk2a1, Olfm1, Atp6v0a1, Arpc2, Lin7a, Chl1, Dlg3, Sptbn4 |
| *transport vesicle membrane* | 21 | Yes (30) | 5.1E-06 | Rab3c, Sv2b, Rab3a, Atp6v0a1, Lin7a, Syngr3, Syngr1, Lin7c, Dnajc5, Map6, Syp, Syn1, Stx1a, Pdcd6, Syt7 |
| *site of DNA damage* | 6 | Yes (4) | 2.3E-04 | Arpc2, Arpc4, Actr3, Actr2, Arpc1a, Cul5 |

*Italics: RRVGO analysis, parentheses indicate number of child GO terms inside cluster

**C**

**Overrepresented SynGO:BP Terms in Downregulated Proteins (WT-CTL vs TG-CTL, 10 moa)**

| Term and No. of Proteins | | P-val | Top 15 Proteins |
|---|---|---|---|
| synaptic vesicle cycle | 25 | 2.0E-06 | Stxbp1, Sv2b, Rab3a, Atp6v0a1, Syngr3, Syngr1, Dnajc5, Ap2m1, Ppfia3, Syp, Rimbp2, Ap2s1, Ap2b1, Abi1, Syn1 |
| process in the presynapse | 26 | 7.0E-05 | Aak1, Stxbp1, Sv2b, Rab3a, Atp6v0a1, Syngr3, Syngr1, Dnajc5, Ap2m1, Ppfia3, Syp, Rimbp2, Ap2s1, Ap2b1, Abi1 |
| synaptic vesicle exocytosis | 12 | 1.5E-04 | Stxbp1, Sv2b, Rab3a, Ppfia3, Syp, Rimbp2, Stx1a, Syt7, Vamp2, Rph3a, Git1, Prkcg |
| synapse organization | 23 | 5.5E-04 | Dlg3, Psd, Ppfia3, Actr3, Rimbp2, Nrcam, Plxna4, Abi1, Dgkz, Mark2, Nckap1, Dlg1, Cap2, Rap2a, Actb |
| regulation of postsynaptic membrane neurotransmitter receptor levels | 14 | 5.5E-04 | Dlg3, Akap5, Ap2m1, Gphn, Ap2s1, Ap2b1, Dlg1, Rap2a, Lgi1, Cntnap2, Kalrn, Git1, Synj1, Hpca |
| postsynaptic neurotransmitter receptor endocytosis | 6 | 8.2E-03 | Akap5, Ap2m1, Ap2s1, Ap2b1, Synj1, Hpca |
| structural constituent of synapse | 6 | 2.8E-02 | Dlg3, Rimbp2, Dlg1, Actb, Git1, Mpp2 |
| structural constituent of postsynapse | 5 | 2.8E-02 | Dlg3, Dlg1, Actb, Git1, Mpp2 |
| regulation of synaptic vesicle cycle | 4 | 2.9E-02 | Syngr3, Syngr1, Dnajc5, Syn1 |
| regulation of synaptic vesicle exocytosis | 4 | 4.5E-02 | Sv2b, Syp, Git1, Prkcg |
| regulation of postsynaptic neurotransmitter receptor activity | 3 | 4.5E-02 | Src, Cnih2, Begain |
| neurotransmitter receptor localization to postsynaptic specialization membrane | 5 | 4.5E-02 | Gphn, Dlg1, Lgi1, Kalrn, Git1 |
| structural constituent of postsynaptic density | 3 | 4.9E-02 | Dlg3, Dlg1, Mpp2 |

**D**

**SynGO:BP analysis of downregulated proteins in 10 moa WT-ES vs TG-ES**

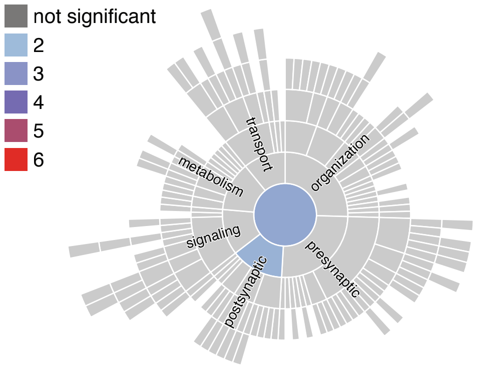


**SynGO:CC analysis of downregulated proteins in 10 moa WT-ES vs TG-ES**

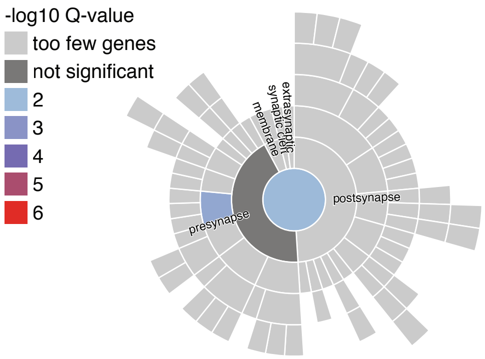


**E**

**Top 5 Overrepresented GO:BP Terms (10 moa WT-CTL vs WT-ES Cohort 3)**

| Term* | Size | RRVGO* | P-val | (Top 15) Proteins |
|---|---|---|---|---|
| negative regulation of blood pressure | 3 | No | 9.0E-03 | Rnpep, Snx5, G6pdx |
| *sarcoplasmic reticulum calcium ion transport* | 2 | Yes (6) | 2.9E-02 | Ryr2, Atp2a2 |
| *regulation of cardiac muscle cell action potential* | 2 | Yes (7) | 2.9E-02 | Ryr2, Atp2a2 |
| *regulation of cardiac muscle contraction by calcium ion signaling* | 2 | Yes (14) | 2.9E-02 | Ryr2, Atp2a2 |
| *regulation of actin filament-based movement* | 2 | Yes (5) | 3.6E-02 | Ryr2, Atp2a2 |
| *left ventricular cardiac muscle tissue morphogenesis* | 1 | Yes (1) | 4.4E-02 | Ryr2 |

*Italics: RRVGO analysis, parentheses indicate number of child GO terms inside cluster

**F**

**Top 5 Overrepresented GO:CC Terms (10 moa WT-CTL vs WT-ES Cohort 3)**

| Term* | Size | RRVGO* | P-val | (Top 15) Proteins |
|---|---|---|---|---|
| cytosol | 10 | No | 4.1E-02 | Atic, Nsfl1c, Snx5, Stk11, G6pdx, Pdxk, Cab39, Flii, Pafah1b3, Fbxo2 |
| centriolar satellite | 2 | No | 4.1E-02 | G6pdx, Flii |
| serine/threonine protein kinase complex | 2 | No | 4.1E-02 | Stk11, Cab39 |
| protein kinase complex | 2 | No | 4.1E-02 | Stk11, Cab39 |
| *sarcoplasmic reticulum membrane* | 2 | Yes (4) | 1.4E-02 | Ryr2, Atp2a2 |
| *endoplasmic reticulum membrane* | 4 | Yes (3) | 1.4E-02 | Ryr2, Atp2a2, Gdpd1, Cds2 |
| *nuclear outer membrane-endoplasmic reticulum membrane* | 4 | Yes (2) | 1.4E-02 | Ryr2, Atp2a2, Gdpd1, Cds2 |
| *calcium ion-transporting ATPase complex* | 1 | Yes (3) | 1.9E-02 | Atp2a2 |
| *extrinsic component of cytoplasmic side of plasma membrane* | 2 | Yes (3) | 2.4E-02 | Ryr2, Atp2a2 |

*Italics: RRVGO analysis, parentheses indicate number of child GO terms inside cluster

**G**

**Top 5 Overrepresented GO:BP Terms (10 moa TG-CTL vs TG-ES Cohort 3)**

| Term* | Size | RRVGO* | P-val | (Top 15) Proteins |
|---|---|---|---|---|
| catabolic process | 157 | Yes (14) | 2.8E-07 | Pkm, Paip1, Aldoc, Pabpc1, Gstm1, Pgk1, Pgd, Pfkm, Snx1, Ube2v2, Pgam1, Tpi1, Aldh1l1, Psmd2, Lhda |
| organonitrogen compound metabolic process | 277 | Yes (18) | 4.5E-06 | Phgdh, Pkm, Paip1, Aldoc, Pabpc1, Gad1, Adk, Acly, Gstm1, Pgp, Pgk1, Aldh7a1, Pfkm, Gpx4, Ppm1a |
| small molecule biosynthetic process | 46 | Yes (6) | 5.1E-04 | Phgdh, Pkm, Gad1, Adk, Acly, Gstm1, Pgp, Pgk1, Pgd, Tpi1, Ldha, Fasn, Ldhb, Got1, Impa1 |
| carbohydrate metabolic process | 45 | Yes (5) | 8.9E-04 | Pkm, Aldoc, Pgp, Pgk1, Pgd, Pfkm, Pgam1, Lancl2, Tpi1, Ldha, Mpi, Pgls, Got1, Impa1, Akr1a1 |
| protein modification by small protein conjugation or removal | 42 | Yes (6) | 1.2E-03 | Ube2v2, Ube2n, Cops4, Uba1, Usp9x, Fbxo2, Trim3, Atg3, Skp1, Usp5, Gclc, Fbxo6, Cops5, Cops7a, Ube2o |
| lipid catabolic process | 4 | No | 4.4E-02 | Hadha, Hadhb, Abhd6, Decr1 |
| fatty acid catabolic process | 3 | No | 4.4E-02 | Hadha, Hadhb, Decr1 |
| fatty acid beta-oxidation | 3 | No | 4.4E-02 | Hadha, Hadhb, Decr1 |
| monocarboxylic acid catabolic process | 3 | No | 4.4E-02 | Hadha, Hadhb, Decr1 |
| organic acid catabolic process | 4 | No | 4.4E-02 | Hadha, Hadhb, Bckdha, Decr1 |

*Italics: RRVGO analysis, parentheses indicate number of child GO terms inside cluster

**H**

**Top 5 Overrepresented GO:CC Terms (10 moa TG-CTL vs TG-ES Cohort 3)**

| Term* | Size | RRVGO* | P-val | (Top 15) Proteins |
|---|---|---|---|---|
| *cytosol* | 302 | Yes (2) | 2.3E-26 | Pkm, Kpnb1, Paip1, Aldoc, Rab6b, Pabpc1, Ykt6, Adk, Acly, Me1, Gstm1, Pgk1, Tbcb, Pgd, Aldh7a1 |
| *nucleus* | 231 | Yes (3) | 1.6E-07 | Pkm, Kpnb1, Pabpc1, Adk, Acly, Aldh7a1, Pfkm, Gpx4, Ppm1a, Ube2v2, Pgam1, Lancl2, Ywhae, Tpi1, Psmd2 |
| *proteasome complex* | 25 | Yes (8) | 3.6E-07 | Psmd2, Psmd12, Psma2, Psma1, Psma7, Psmd3, Psmb1, Psmc2, Txnl1, Psmb5, Psma6, Psmd8, Psmd13, Psmb4, Psmb7 |
| *microtubule* | 49 | Yes (5) | 1.2E-05 | Tbcb, Dnm3, Klc1, Pafah1b1, Kif5c, Map6, Dnm1, Dctn1, Kif5b, Synj1, Arfgef2, Kif21a, Dnm1l, Kif2a, Dync1i2 |
| *microtubule cytoskeleton* | 79 | Yes (6) | 1.2E-04 | Tbcb, Dctn4, Mdh1, Dnm3, Ywhae, Klc1, Pafah1b1, Kif5c, Map6, Mpp1, Dnm1, Dctn1, Pdcd6ip, Brsk2, Psma1 |
| mitochondrial fatty acid beta-oxidation multienzyme complex | 2 | No | 1.0E-03 | Hadha, Hadhb |
| fatty acid beta-oxidation multienzyme complex | 2 | No | 1.0E-03 | Hadha, Hadhb |

*Italics: RRVGO analysis, parentheses indicate number of child GO terms inside cluster
