## Supplementary figures and images for "Early-life stress exposure impacts the hippocampal synaptic proteome in a mouse model of Alzheimer’s disease: age- and pathology-dependent effects on mitochondrial proteins"

### 20230301_Figure_S1_redo.pdf

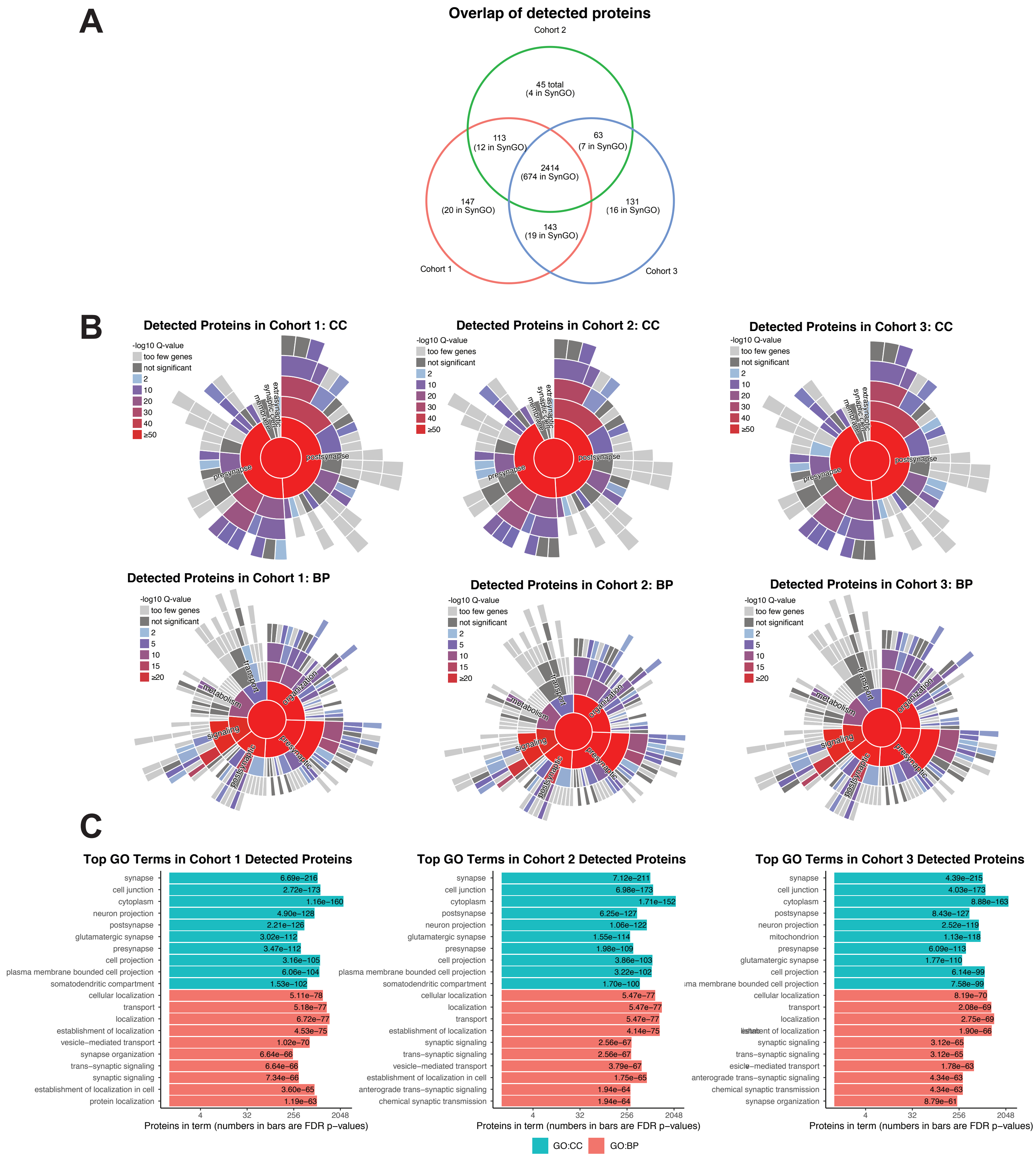

### 20230301_Figure_S3_redo-compressed.pdf

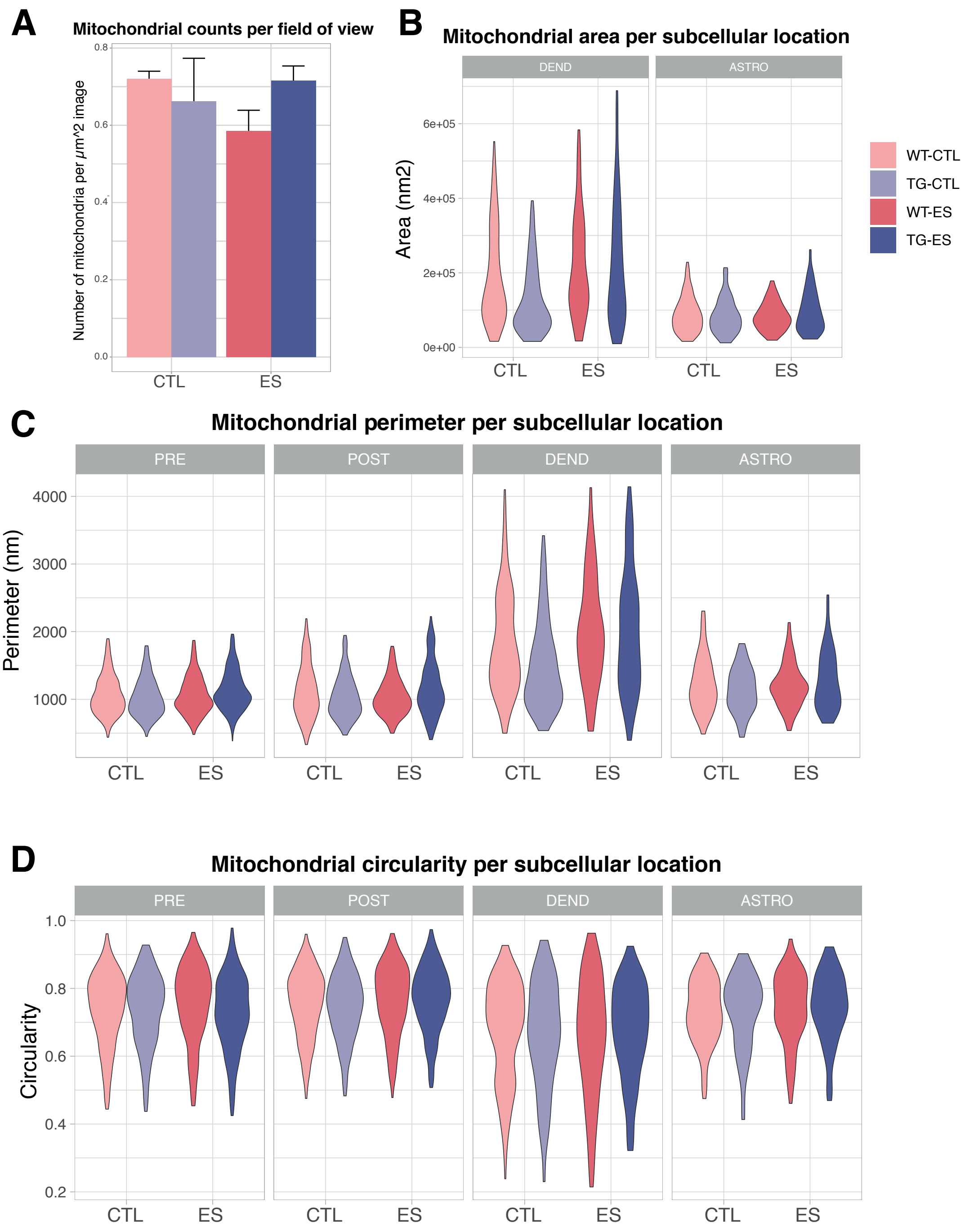
